# Functional relevance of CASP16 nucleic acid predictions as evaluated by structure providers

**DOI:** 10.1101/2025.04.15.649049

**Authors:** Rachael C. Kretsch, Reinhard Albrecht, Ebbe S. Andersen, Hsuan-Ai Chen, Wah Chiu, Rhiju Das, Jeanine G. Gezelle, Marcus D. Hartmann, Claudia Höbartner, Yimin Hu, Shekhar Jadhav, Philip E. Johnson, Christopher P. Jones, Deepak Koirala, Emil L. Kristoffersen, Eric Largy, Anna Lewicka, Cameron D. Mackereth, Marco Marcia, Michela Nigro, Manju Ojha, Joseph A. Piccirilli, Phoebe A. Rice, Heewhan Shin, Anna-Lena Steckelberg, Zhaoming Su, Yoshita Srivastava, Liu Wang, Yuan Wu, Jiahao Xie, Nikolaj H. Zwergius, John Moult, Andriy Kryshtafovych

## Abstract

Accurate biomolecular structure prediction enables the prediction of mutational effects, the speculation of function based on predicted structural homology, the analysis of ligand binding modes, experimental model building and many other applications. Such algorithms to predict essential functional and structural features remain out of reach for biomolecular. Here, we report quantitative and qualitative evaluation of nucleic acid structures for the CASP16 blind prediction challenge by 12 of the experimental groups who provided nucleic acid targets. Blind predictions accurately model secondary structure and some aspects of tertiary structure, including reasonable global folds for some complex RNAs, however, predictions often lack accuracy in the regions of highest functional importance. All models have inaccuracies in non-canonical regions where, e.g., the nucleic-acid backbone bends or a base forms a non-standard hydrogen bond. These bends and non-canonical interactions are integral to form functionally important regions such as RNA enzymatic active sites. Additionally, the modeling of conserved and functional interfaces between nucleic acids and ligands, proteins, or other nucleic acids remains poor. For some targets, the experimental structures may not represent the only structure the biomolecular complex occupies in solution or in its functional life-cycle, posing a future challenge for the community.

## 1 Introduction

Structural biologists have realized the immense benefits of accurate computational structure prediction methods now available for proteins^1^, but the accuracy of computational nucleic acid structure prediction lags far behind^2^,CASP16 NA assessment paper. The scientists who determine experimental structures play an integral role in the field’s journey towards accurate structure prediction. The structures they determine increase the amount of data from which prediction algorithms can learn and enable blind structure prediction challenges like the Critical Assessment of Techniques for Structure Prediction (CASP)CASP16 NA assessment paper and RNA puzzles^3^. Additionally, nucleic acid biologists and experimental structural biologists identify the features of nucleic acid structure that are important to predict accurately for functional or biophysical reasons, guiding the goals of structure prediction. In this manuscript, as in CASP15^4^, structure providers for the CASP16 experiment explain important structural and functional features of their nucleic acid-containing structures and analyze how accurately modelers predicted these features.

In the second iteration of the nucleic acid category of CASP (CASP16, 2024), 37 RNA, 1 DNA, 11 RNA-multimer, 16 nucleic-acid-protein complexes, and 6 nucleic-acid ligand complexes were provided as modeling targets by 22 structure determination groups from 10 countries. All targets were released for prediction from May to July 2024. Among these, 12 targets were solved by X-ray crystallography, 29 by cryogenic electron microscopy (cryo-EM), and 1 by nuclear magnetic resonance (NMR).

All target providers were invited to contribute to this paper, resulting in 12 sections highlighting 18 of the targets (**Table 1**). The numerical evaluation of CASP16 models is available at the Prediction Center website (see **Data Availability** statement). A general evaluation of predicted accuracy over all targets, are provided elsewhere in this issueCASP16 NA assessment paper.

**Table 1:**
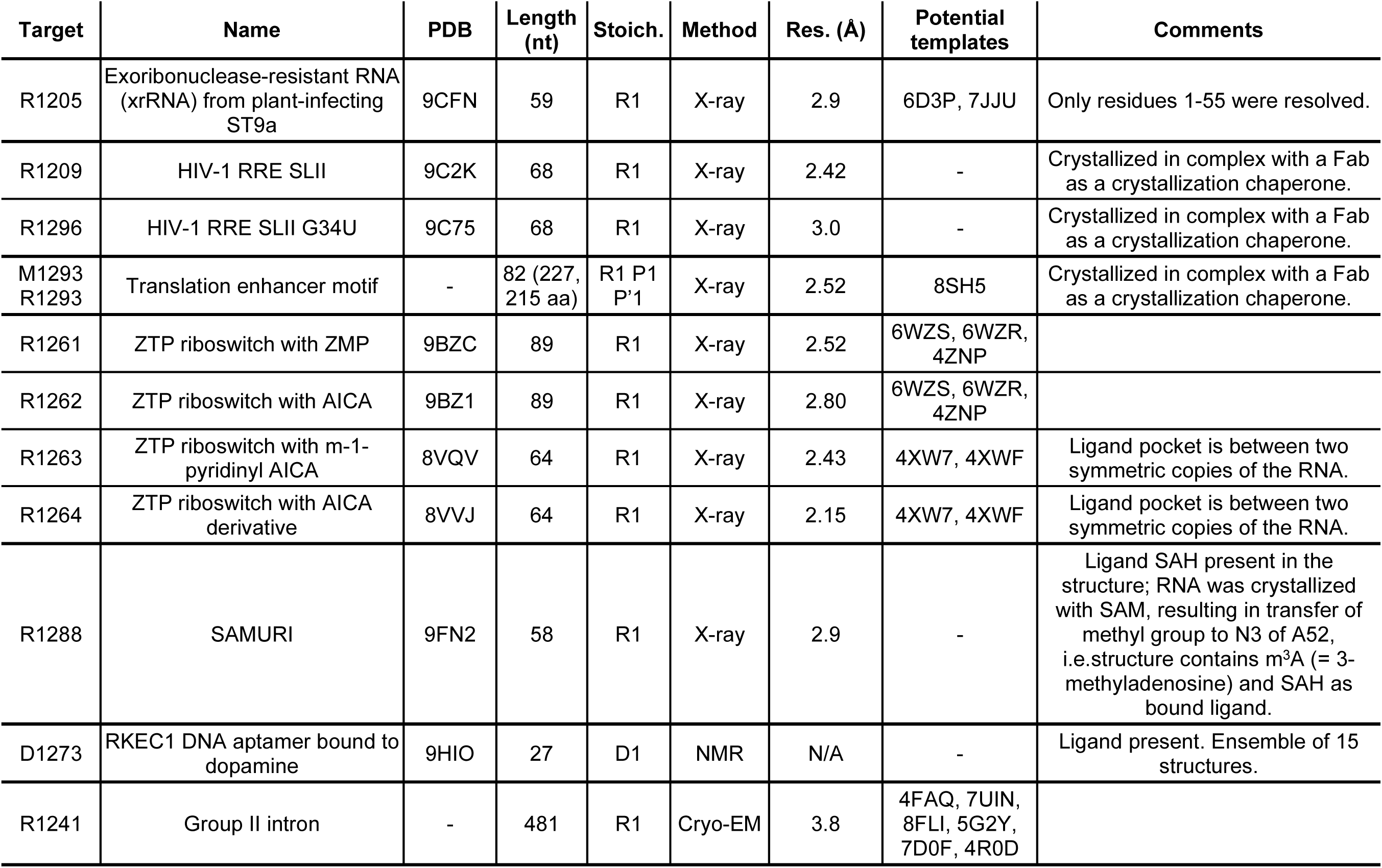
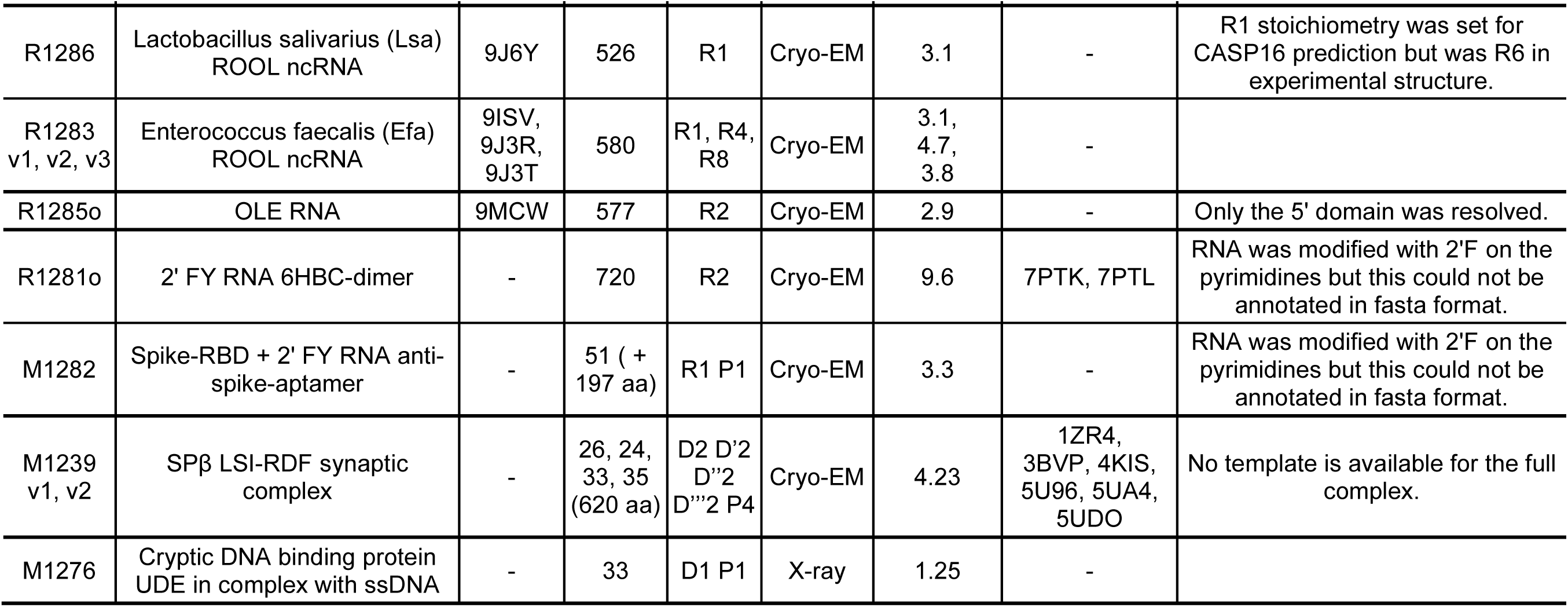
CASP16 RNA targets included in this study.

## 2 Results

### 2.1 Exoribonuclease-resistant viral RNA (CASP: R1205, PDB: 9CFN) provided by Jeanine Gezelle and Anna-Lena Steckelberg

Exoribonuclease-resistant RNAs (xrRNAs) are specialized viral RNA structures that block degradation by cellular 5′-3′ exoribonucleases at specific sites within the viral RNA genome^5^. This resistance results in the accumulation of viral subgenomic RNAs, which play critical roles in immune modulation and viral replication^6^. Initially identified in human-pathogenic flaviviruses, such as Dengue virus and Zika virus, xrRNAs have since been found across a wide range of RNA virus families infecting both animals and plants^7–10^. Despite a complete lack of sequence similarity, xrRNA from distantly related virus families share a conserved core motif: a central ring encircling the 5′ end of the RNA^5,8,11^. This ring functions as a molecular brace, effectively countering the unidirectional unwinding forces of cellular exoribonucleases to protect the RNA from degradation.

CASP16 target R1205 (PDB: 9CFN) represents the first high-resolution structure of a plant-virus xrRNA in its nuclease-resistant conformation^12^. The structure, derived from the plant-infecting ST9a virus, was solved by X-ray crystallography to 2.9 Å resolution. The crystallization construct consisted of 59 nucleotides, with nucleotides 56-59 excluded from the final model due to insufficient electron density at the 3′ end. The RNA adopts a compact fold composed of a stem loop (SL) of coaxially stacked helices P1 and P2, with a 9-base pair pseudoknot (PK) between the apical loop and the 3′ end of the RNA (**Figure 1A-B**). The PK causes the 3′ end to wrap around the 5′ end, forming the characteristic protective ring topology previously observed in xrRNAs from other viral families (**Figure 1B**). A highly conserved network of non-canonical interactions orients the helices and anchors the ring within the xrRNA scaffold (**Figure 1A-B**). Most notably, a hairpin loop, L2B, protrudes from the apical loop L2 to cap the base of the PK. The backbone of L2B resembles a U-turn RNA tetraloop, but it is closed by a reverse Hoogsteen U-A base pair and a reverse Watson-Crick base pair involving flipped-out A31 of P2. L2B is positioned at an acute angle next to P2, behind the protective ring, and engages in multiple non-canonical tertiary interactions with the J1/2 and J1/3 regions of the RNA. These include (a) A5 stacking between A28 and A29 and interacting with the sugar edge of G26, (b) C6 forming a long-range base pair with G27, and (c) two conserved base triple interactions, A42-A5-G26 and G43-A29-U25, that position the 3′ end of the protective ring. While R1205 represents the first structure of a plant-virus xrRNA in its active PK conformation, two previously solved structures captured related xrRNA variants in a SL conformation (PDB: 6D3P, 7JJU)^8,13^. In these cases, the intramolecular PK was replaced by equivalent trans interactions between two adjacent RNA molecules, forming a ‘domain-swapped’ dimer. These earlier structures could have informed the CASP16 predictions.

**Figure 1.**
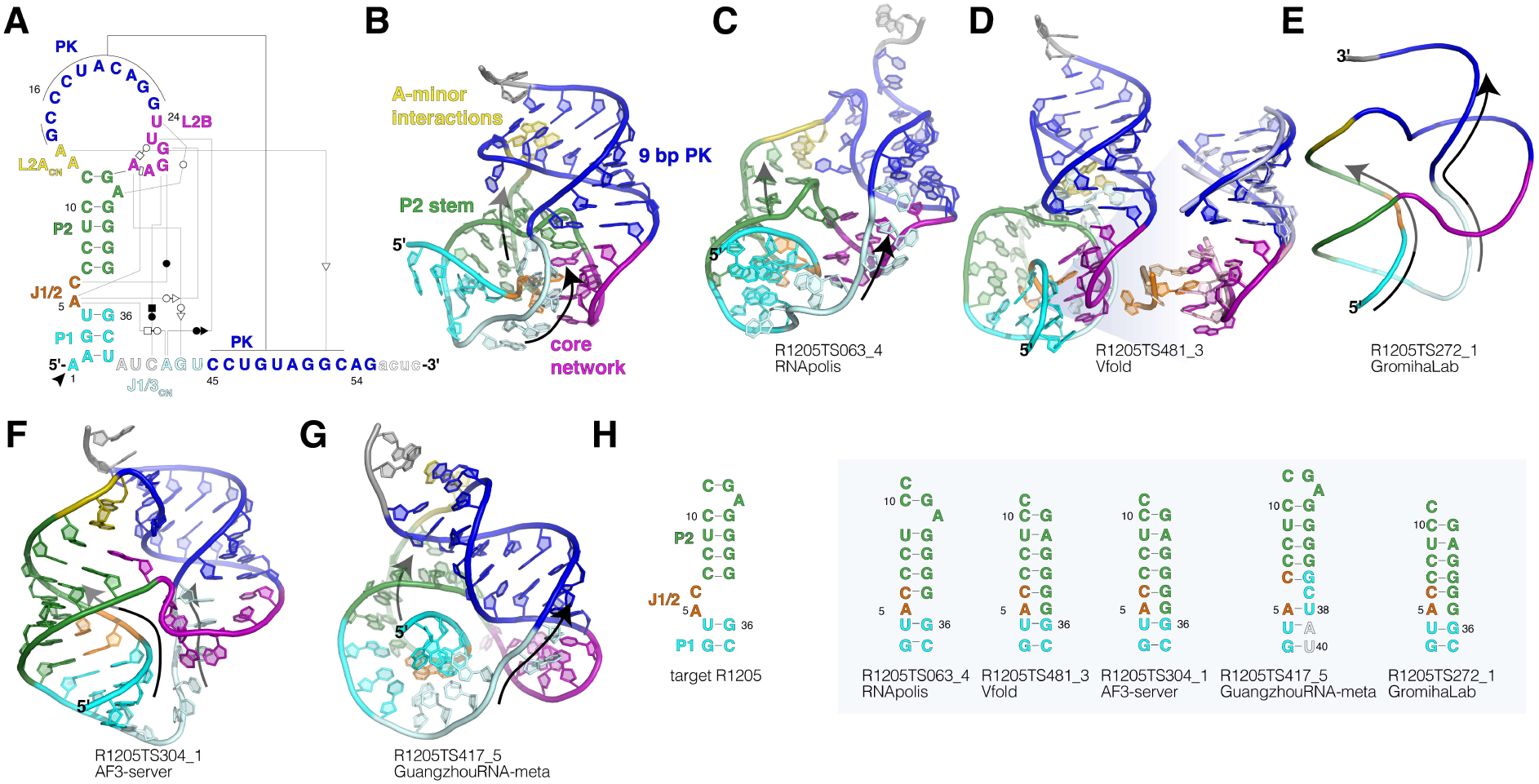
Predictions of the ST9a Xrn1-resistant RNA (CASP: R1205, PDB: 9CFN). (**A**) Secondary structure diagram of ST9a xrRNA crystallization construct. Non–Watson–Crick base pairs are in Leontis–Westhof annotation and the Xrn1 stop site is depicted by the arrow. Non-modeled nucleotides are shown as lowercase letters. (**B**) Ribbon diagram of the ST9a xrRNA crystal structure. Colors match D. Arrows in all panels indicate strand directionality 5′-3′. (**C**) 3D ribbon diagram of the RNApolis group CASP16 prediction with leading metrics, colors to match A. (**D**) Left: 3D ribbon diagram of Vfold prediction TS481_3, colors to match A. Right: overlay of the PK and J1/2-L2B region of the Vfold model (dark) and the experimental crystal structure (PDB: 9CFN, light/tints); RMSD = 1.537. All 3D structure analysis was performed in PyMOL. (**E**) Backbone depiction of the GromihaLab prediction, colors to match A and B, arrows in the 5′-3′ direction. (**F-G**) 3D ribbon diagrams of CASP16 predictions with leading metrics from F) AF3-server and G) GuangzhouRNA-meta, colors to match A. Note that the RNA backbones in D-F form knots. (**H**) Secondary structure depictions of the P1 and P2 regions in the crystal structure (left) and in CASP16 predictions (right).

Many models correctly predicted some aspects of the global fold of R1205—including helices P1 and P2, the PK, and the 3′ end wrapping around the 5′ end—yet all failed to accurately predict crucial non-canonical tertiary interactions (**Figure 1C-G**). The model with the lowest RMSD (7.46 Å) was TS063_4, submitted by the RNApolis group. Surprisingly, TS063_4 and related models from RNApolis (all with an RMSD < 10 Å) were the only structures to correctly predict the L2B hairpin positioned behind the protective ring at the base of the PK (**Figure 1C**). However, key non-canonical interactions of L2B, including the conserved Hoogsteen base pair and base-triple interactions, were missing. Furthermore, the models predicted a shorter 6-base pair PK, highlighting the challenge of folding a structure that accommodates both P1-P2 and the full 9-base pair PK. In the crystal structure, this is achieved by a sharp bend in the L2 backbone, along with two adenosines that interact with the minor groove of the PK **(Figure 1B)**. These unusual A-minor interactions in cis are not found in any other structures currently available in the PDB and were not predicted by any CASP16 models.

The best model according to most CASP metrics was V-fold TS481_3 (GTD-TS 0.43, LDDT 0.499) (**Figure 1D**). This model was the only one to successfully predict a subset of the non-canonical L2B interactions, including the A-U Hoogsteen base pair, A5 stacking between A28 and A29, and the interaction of A5 with the sugar edge of G26 (**Figure 1D**, right). However, while these local interactions were correctly predicted, their position within the global structure was incorrect. Specifically, in this model, L2B is positioned in front of the protective ring. This orientation facilitates the accommodation of both P1-P2 and an 8-base pair PK in the same structure, but it introduces a remarkable folding problem: the resulting RNA structure contains two intertwined helices that form a genuine knot. To form this knot, the 3′ end would need to pass through a closed ring in the RNA, a highly improbable folding path. Notably, we found that a majority of high-scoring models adopted similar knot topologies.

While the xrRNA crystal structure exhibits numerous sharp backbone bends and non-canonical interactions (**Figure 1A-B**), most CASP16 models rely predominantly on canonical A-form helices. This is particularly evident in the AlphaFold3 models, as well those from GromihaLab, KiharaLab, and the GuangzhouRNA group (**Figure 1E-G**). Although these models scored reasonably well in CASP metrics, they all failed to capture key non-canonical interactions. Instead, all created a knot in the RNA backbone, similar to the Vfold model (**Figure 1E-G**). Beyond missing critical tertiary interactions, most models also mispredicted elements of the secondary structure. This is especially evident in P2, where a register shift was introduced due to failure to position C6 and/or A31 in bulged regions engaged in long-range tertiary interactions (**Figure 1H**).

In summary, we found that an overreliance on canonical A-form RNA helices limited the accuracy of most CASP16 models. Despite two related RNA structures being available in the PDB^8,13^, and L2B-like motifs identified in ribosomal RNAs^8^, only one model accurately predicted the L2B tertiary interactions (**Figure 1D**). Additionally, predictions often misinterpreted nucleotides involved in long-range tertiary interactions as part of extended helices (**Figure 1H**). To improve prediction accuracy, more diverse training sets are needed, including additional high-resolution structures of RNA featuring non-canonical interactions. Most remarkably, to maximize base-pairing interactions, many models predicted implausible structures that introduced genuine knots in the RNA backbone (**Figure 1D-G**). Considering folding intermediates in RNA structure predictions could help avoid such issues and better reflect biologically relevant structures.

### 2.2 HIV-1 rev response element stem-loop II (CASP: R1209 and R1296, PDB: 9C2K and 9C75) provided by Manju Ojha and Deepak Koirala

The Rev Response Element (RRE) is a ∼350-nucleotide-long *cis-acting* RNA domain found in the env coding region of the HIV-1 genome^14,15^. RRE plays a crucial role in HIV-1 replication by interacting with the viral-encoded protein Rev to form an oligomeric complex that facilitates the nuclear export of intron-containing viral RNAs during the late phase of viral infection^16–18^. This ribonucleoprotein complex has been considered a promising therapeutic target for developing drugs against HIV infections^19,20^; however, the mechanism underlying this crucial virological process remains poorly understood, mainly due to the lack of high-resolution structural information on RRE and RRE-Rev complexes. The proposed secondary structures of HIV-1 RRE consist of four stem-loop structures II-V budding from a basal stem I^15,21^. Some previous studies based on small-angle X-ray scattering (SAXS) and atomic force microscopy (AFM) proposed an unusual A-shaped tertiary structure of the intact RRE^22,23^, but detailed structural features were missing due to the low resolution of those methods. Recently, Tipo et al^24^ solved the 2.85 Å crystal structure of the stem-loop II (SLII) using a t-RNA scaffold approach (CASP: R1203, PDB: 8UO6) and found that two molecules within the crystallographic unit folded differently, which they called ‘compact’ and ‘extended’ conformations^24^.

Using a Fab-assisted crystallography approach^25,26^, we crystallized and determined the structure of a 68-nucleotide SLII construct (CASP: R1209, PDB: 9C2K, 2.42 Å resolution), where a Fab binding sequence replaced the IIc loop^27^. Our SLII crystal structure revealed a unique three-way junction (3WJ) architecture, where the base IIa stem bifurcates into the IIb and IIc stem-loop structures, with extensive interactions between the junction nucleotides^27^. Interestingly, while the structures of individual domains IIa, IIb and IIc were similar to that determined by Tipo et al.^24^, our structure showed a completely different arrangement within the 3WJ, resulting in a different 3D architecture of the RNA. We also solved the crystal structure of the SLII G34U mutant (CASP: R1296, PDB: 9C75, 3.0 Å resolution), which adopted a different conformation than its non-mutant form, where the junction nucleotides reregistered to form simpler 3WJ^27^. Notably, the non-mutant form appeared more compact than the G34U mutant. Our functional studies have shown that the compact non-mutant fold represents a functional conformation, and the extended G34U mutant induces loss of nuclear export function^27^. Due to the highly plastic nature of the SLII structure and its relation with functional consequences, understanding the multiple possible folds of SLII RNA and how it performs with RNA 3D structure prediction were quite interesting; therefore, crystallized sequences of both non-mutant and G34U-mutant were submitted to the CASP16 for blind predictions.

Many groups that participated in the CASP16 predicted a total of 213 and 230 models of SLII (R1209) and SLIIG34U (R1296). For SLII (R1209), the alignment of crystal structure with the predicted models showed a high deviation, with the top-five global RMSDs ranging from 7.8 Å to 9.4 Å (**Figure 2A-E**). However, with a few minor variations, all top-five predicted models showed a good agreement with the individual subdomain structures: IIa (RMSD = 0.856 ± 0.213 Å; mean ± s.d.), IIb (RMSD = 2.899 ± 0.284 Å; **Figure 2F**), and IIc (RMSD = 0.786 ± 0.230 Å; **Figure 2G**).

**Figure 2.**
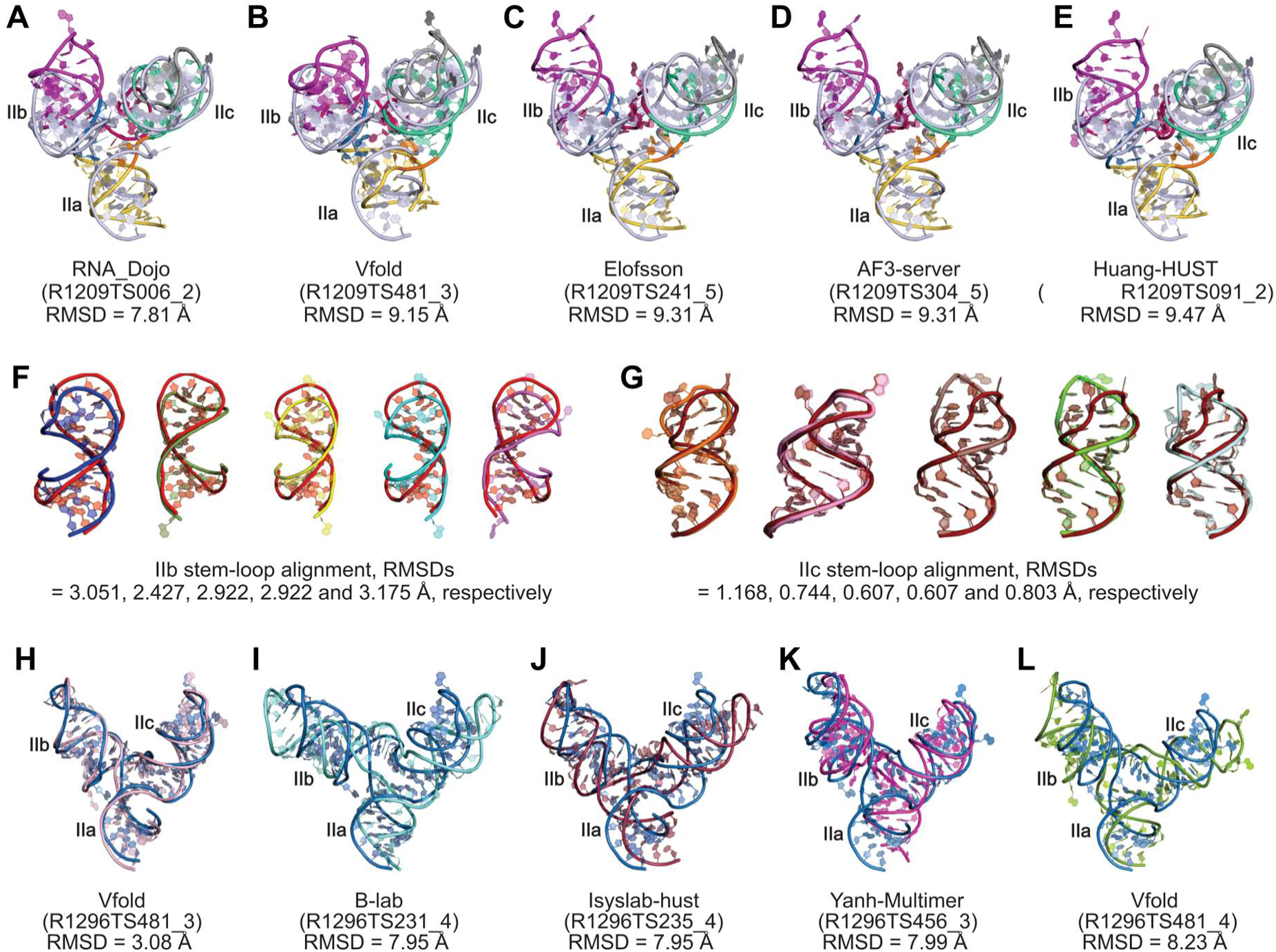
Comparison of top-five predicted models with HIV-1 RRE SLII and SLIIG34U mutant crystal structures (CASP: R1209 and R1296, PDB: 9C2K and 9C75). (**A-E**) The top-five predicted models for SLII (R1209, colored) scored according to the CASP16 global RMSDs and superimposed with the corresponding crystal structure (colored light blue). The alignment of the IIb stem-loops (**F**) and IIc stem-loops (**G**) from the same top-five models with the corresponding regions of the crystal structure. The IIb and IIc stem-loop models are colored differently, but corresponding crystal structures are colored the same for facile comparisons. The RMSDs shown in (**F**) and (**G**) represent all-atom alignment in PyMol. (**H-L**) The top-five predicted models for SLIIG34U (R1296, colored differently) scored according to the CASP16 global RMSDs and superimposed with the corresponding crystal structure (colored blue).

The IIb (**Figure 2F**) showed higher variability than IIc (**Figure 2G**) for all five models, which seems more dynamic, as indicated by high crystallographic B factors for this region. These comparisons, therefore, show that the significant discrepancies between the crystal structure and predicted models originate from the 3WJ region that defines the relative orientations of these stem-loop structures. Even for the model R1209TS006_2 predicted by RNA_Dojo with the lowest global RMSD (7.81 Å), the base-pairing and tertiary interaction patterns within the 3WJ are quite different from those observed in the crystal structure. For example, in the predicted model, the Jab G9 makes a non-canonical pairing with Jca U64, but in the crystal structure, it makes a tertiary interaction with C32, and U64 remains unpaired. G13 is base-paired with C32 in the model but forms a non-canonical G13•A31 pair in the crystal structure. For all top-five models, A31 has been predicted to be flipped out as a bulge. Except for the model R1209TS006_2 and the crystal structure, all other top-four models showed Jca A63 and U64 base-pairing with the G40 and C41, respectively.

Compared to SLII (R1209), the predicted models for SLIIG34U (R1296) aligned better with the corresponding crystal structure, with the top-five global RMSDs ranging from 3.08 Å to 8.23 Å (**Figure 2H-L**). Similar to the R1209, all top-five predicted models showed high resemblance for individual subdomains: IIa (RMSD = 0.771 ± 0.075 Å), IIb (RMSD = 1.339 ± 0.158 Å), and IIc (RMSD = 0.566 ± 0.220 Å), suggesting that the variations between the crystal structure and predicted models reflects the discrepancies within the 3WJ region. Nevertheless, unlike R1209, one of the models, R1296TS481_3 predicted by Vfold, is almost identical to the crystal structure with a global RMSD of 3.08 Å. The only noticeable difference within the 3WJ is the wobble pairing between Jbc G40 and Jca U64 in the model versus no pairing in the crystal structure, even though the position and helical stacking of these nucleotides are similar. Because the 3WJ of SLIIG34U is relatively simple with no evident tertiary interactions than those in non-mutant SLII, the blind predictions perhaps performed better for the mutant sequence.

Our experimental SLII and SLIIG34U structures captured in crystals demonstrate how a point mutation in an RNA structure can drastically alter its overall conformation. However, solving such conformationally diverse structures is often not feasible due to the challenges associated with RNA structural determination. Therefore, the high similarity of the predicted model and SLIIG34U highlights the value of blind prediction computational approaches for understanding 3D folding and the complex conformational landscape that can emerge from an RNA sequence in solution. However, such an RNA structural plasticity complicates the prediction of functionally relevant 3D structures. In these two cases (R1209 and R1296), whereas the predicted structures of subdomains for most models align closely with the corresponding crystal structures, the relative positioning of these subdomains seems to be the key factor determining the divergence between the experimental and predicted results. Because crystal structures only provide a static 3D reference for comparisons with predicted models, it remains unclear whether the deviations between the crystal structure and predicted models (in-solution or near-native state) originate from the relative orientations of subdomains within the RNA structure or whether the crystal structure itself is limited to a specific fold due to the stabilization by lattice contacts. For instance, the crystal structure of SLII showed significant crystal contacts between the symmetry-mate nucleotides, whereas SLIIG34U exhibited no apparent crystal contacts near 3WJ. While blind prediction of RNA 3D structures has improved significantly in recent years, acquiring more high-resolution experimental structures using various methods (X-ray, NMR and cryo-EM) helps advance the RNA structural database that, in turn, enriches the computational algorithms for better prediction.

### 2.3 3′ cap-independent translation enhancer (3′ CITE) from thin paspalum asymptomatic virus (CASP: R1293/M1293) provided by Anna Lewicka, Yoshita Srivastava and Joseph A. Piccirilli

In eukaryotes, processed messenger RNA (mRNA) is covalently modified with a 5′ N7-methylguanine triphosphate cap and 3′ poly(A) tail, which serve to identify the mRNA by the eukaryotic translation initiation factors^28^. Recognition of the 5′ cap structure by eukaryotic initiation factor 4E (eIF4E) represents a hallmark feature in the initiation of protein translation.

Some RNAs that lack a 5′-cap have evolved mechanisms to circumvent this canonical mechanism. For example, positive strand RNA viruses that replicate in the cytoplasm cannot access the cellular capping machinery present in the nucleus. To enable the virus to exploit the host’s translation machinery, the 5′ cap is functionally replaced with an RNA sequence that folds into a structured element^29^. Recently discovered cap-independent translation enhancers (CITEs) have been found in plant viruses. These structured RNA translation elements located within the 3′ UTR, usually span 100–200 nt in length and encompass a single domain. Their complex tertiary structures are capable of binding host factors that facilitate translation of viral genes^30^.

There are seven classes of CITEs characterized based on their RNA structures and functional identity^31^. One of the classes, named PTE (PMV or PEMV-like translational enhancer) binds to eIF4E using an T-shaped structured RNA element, which contains a G-rich region (G domain, J 2/3) that exhibits apparent complementarity to the C-rich region (C domain, J4/5) located in a three-way between P4 and P5 (**Figure 3A-B**)^32,33^. The crystal structure of the PTE from Pea enation mosaic virus 2 (PEMV2) RNA in complex with structural chaperone, Fab BL3–6, revealed that the G-rich and C-rich regions interact through a complex network of long-range interactions^32^. The long-range interactions between the G domain, C domain and the J5/3 form an unusual architecture that involves parallel strand WCF base pairing within a unique stacked four-layer scaffold that projects a single G of the G domain away from the core fold into solvent, where it is poised to engage in eIF4E binding (**Figure 3A-C**)^33^.

**Figure 3.**
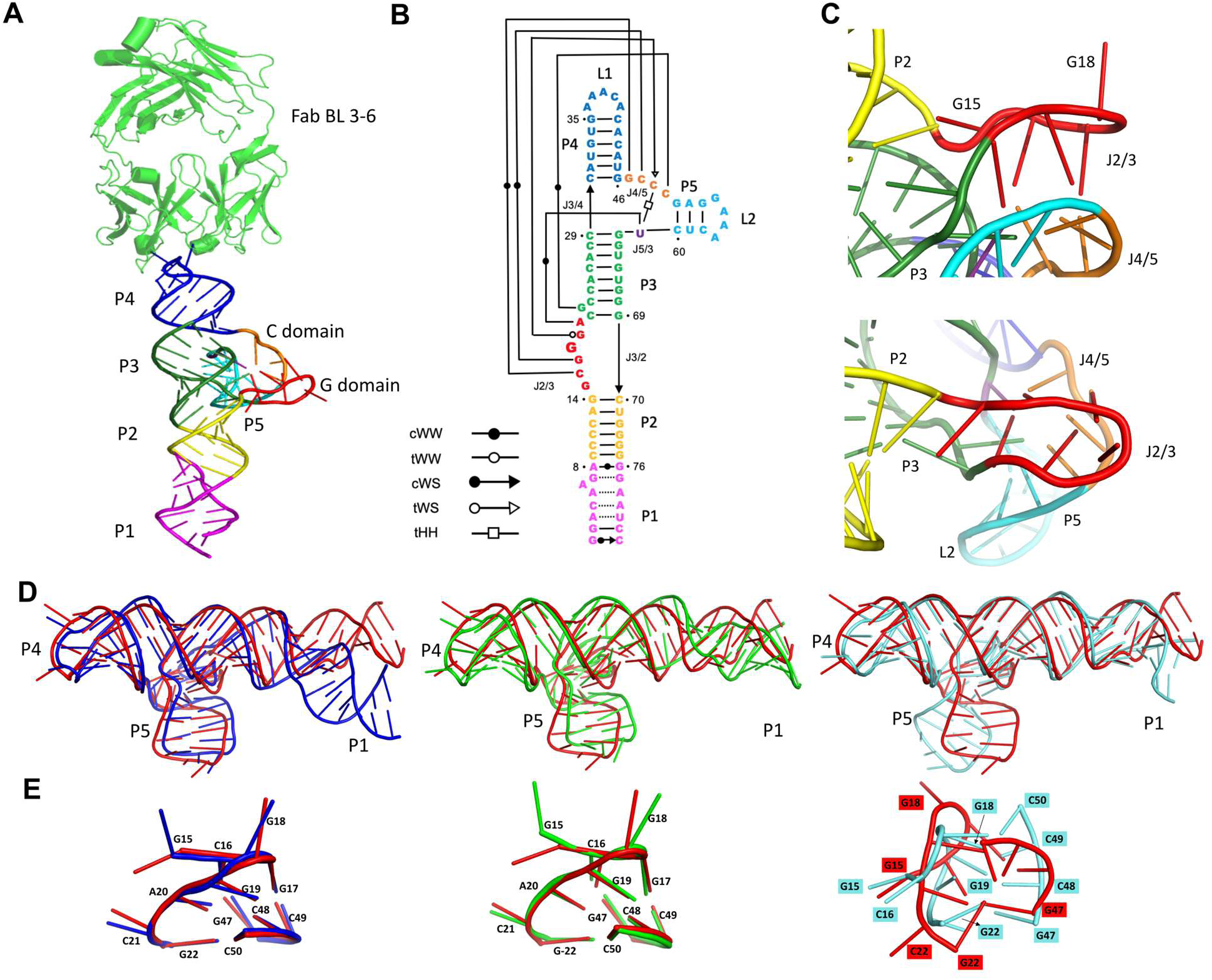
Prediction accuracy for RNA CITE element (CASP: R1293/M1293). (**A**) Crystal structure of TPAV in complex with Fab BL3-6. (**B**) Secondary structure diagram of TPAV RNA (**C**) G and C domains of TPAV (**D**) From left to right: Superposition of Vfold model (R1293TS481_1, blue) on TPAV RNA (red), Superposition of RNApolis model 2 (R1293TS063_2, green) on TPAV RNA, Superposition of GeneSilico model (R1293TS338_1, cyan) on TPAV RNA. E) From left to right: Superposition of paired G and C domains of Vfold model (R1293TS481_1) on TPAV, Superposition of paired G and C domains of RNApolis model (R1293TS063_2) on TPAV, Superposition of paired G and C domains of GeneSilico model (R1293TS338_1) on TPAV.

Recently, we have crystallized and solved the structure of another PTE class homolog from thin paspalum asymptomatic virus (TPAV). The TPAV CITE differs slightly from PEMV2 in its G and C domains. To obtain diffraction-quality crystals, the solvent-exposed RNA loop L1, was mutated to the Fab-binding sequence (GAAACAC).

We submitted the structure of this Fab-RNA complex to CASP16 for structure prediction and received over 200 models for the RNA and over 150 models for the RNA-Fab hybrid. We analysed the models by RMSD and global metrics such as Global Distance Test-Total Score (GDT-TS). We chose to further analyse the models with less than 10 Å RMSD. These models also had a better Local Distance Difference Test (LDDT) and GDT-TS scores. We noticed that most models within the specified RMSD cutoff were able to predict the overall fold of the RNA reasonably well. Additionally, we also looked at the INF (Interaction Network Fidelity) scores and noticed that many models correctly predict the secondary structure of the RNA. With the structure of homologous PEMV2 being available, we anticipated that prediction efforts might utilize it as a modelling template. Some of the models that stood out for overall agreement with the crystal structure were from RNApolis (R1293TS063_2), Vfold (R1293TS481_1) and GeneSilico groups (R1293TS338_1) (**Figure 3D**).

One of the major challenges for modelling was the interaction between the G domain in the asymmetric bulge between P2 and P3 and the C domain between P4 and P5. The G and C domains in our TPAV structure make long range WC interactions to form a conformation suitable for eIF4E binding. In the crystal structure, the G domain makes a bend at position 17G, which changes the direction of RNA. This abrupt turn flips the G18 nucleobase out of the stacked core towards the solvent. The backbone reversal due to the acute turn at G18 places G19 and A20 below G17 and C16, respectively, where they make interactions with C49 and U61, the single J5/3 nucleotide. Upon inspecting the models closely, we found that some of the best models based on global metrics deviated from the crystal structure in the paired interactions between the G and C domains. RNApolis and Vfold do a relatively better job than GeneSilico at predicting the paired G and C domains with the regional RMSD < 2 Å. A bottleneck in modelling these long-range interactions seems to be in modelling of bends and turns in the RNA sugar backbone. The model from GeneSilico tries to constrain the G-C domain to form a more commonly observed anti-parallel helix structure. A specific difference between the predicted models and the crystal structure is the conformation of the G domain’s first residue (G15). In the crystal structure this residue inserts into the pocket created between the P2-P3 stack on one side and the interacting G and C domains on the other side. The nucleobase of G15 is almost perpendicular to the plane of the A13.U71 base pair. G15 forms phosphate-ribose zipper interactions with A20, similar to A18 and A23 in PEMV2, but with an additional interaction between G15:N2 and A20:O3’. In the recently reported structure of the PTE from saguaro cactus virus^26^ the first residue of G domain adopts a similar orientation as observed here for TPAV. In contrast, the predictions for the TPAV element model this residue as being flipped out from the core into solvent. The difference between the crystal structure and the predictions may result from modelling the structure against the published PEMV2 structure, where the corresponding nucleotide is flipped out of the core.

A general observation from the predictions was that most models do a good job describing base-paired regions but predict bulge and junction regions less reliably. This shortcoming shows up in the R1293TS481_1 modelled by Vfold, in which A6 in the core changes the angle between P1 and P2 (**Figure 3B**). This outcome probably reflects model bias towards maintaining a strictly A form helical structure in RNA.

We also analyzed the predictions for the RNA-protein complex. For the complex, we mainly focused on the interface region. We noticed that most models with high global QS metric were able to accurately predict the RNA-protein interfaces. This outcome is anticipated as the RNA-protein interface is well established in literature due to the Fab and its antigenic loop serving as a portable crystallization module. RNApolis, which predicted one of the best models for the RNA component, also predicts the RNA-protein interface quite accurately with an RMSD of 1 Å and a global QS score of 0.96.

### 2.4 ZTP riboswitches bound to synthetic and natural ligands (CASP: R1261, R1262, R1263, and R1264, PDB: 9BZC, 9BZ1, 8VQV, and 8VVJ) provided by Christopher P. Jones

The targets R1261, R1262, R1263, and R1264 are co-crystal structures of ZTP (5-aminoimidazole-4-carboxamide riboside 5’-triphosphate) riboswitches bound to various ligands. ZTP riboswitches often regulate genes associated with purine biosynthesis and folate metabolism^34^ by sensing the levels of ZMP (5-aminoimidazole-4-carboxamide riboside 5’-phosphate) and ZTP. Their conserved folds are characterized by four paired elements that include one pseudoknot. However, variations on this conserved fold include the insertion of additional paired elements.

R1261 and R1262 contain novel structures of the *Clostridium beijerinckii* ZTP riboswitch co-crystallized with ZMP and AICA, respectively. R1261 and R1262 differ in that R1261 was treated with CsCl (for phasing), which may explain an alternative loop conformation present near the Cs^+^ binding site. The *C. beijerinckii* ZTP riboswitch contains an additional paired element inserted between the two conserved sub-domains^35^, and the RNA is relatively modular compared to other ZTP riboswitches, making it an attractive specimen for folding studies^36,37^. As these structures are related to those of previously determined ZTP riboswitches, their prediction should be low-hanging fruit. Indeed, predictions for this RNA were generally acceptable with 72% and 77% of predictions achieving TM-score greater than 0.45 for R1261 and R1262 respectively. However, predictors were not generally able to achieve atomic accuracy (TM-score >0.70): only 4 models (1.8%) for R1261 and 1 (0.5%) for R1262 achieved a TM-score greater than 0.70. The RNA-ligand interaction was predicted accurately, as expected because of the availability of a template: 45% and 31% of predictions for R1261 and R1262 respectively achieved a lDDT for the RNA-ligand interface greater than 0.75.

R1263 and R1264 are co-crystal structures of two synthetic ligands with the previously solved *Schaalia odontolytica* ZTP riboswitch^38^. These synthetic ligands were developed as part of an ongoing investigation of the types of ligands that bind and activate the ZTP riboswitch *in vitro* and *in vivo*^39^. R1263 contains the best synthetic ligand, *m*-1-pyridinyl-AICA, determined at a higher resolution than previously achieved^39,40^ and should be easily predicted. 33% and 26% of the models submitted for R1263 and R1264 respectively were of a high quality (TM-score > 0.70), more than for *C. beijerinckii* ZTP riboswitch, likely because of the better RNA template. R1264 contains a novel ligand (1,8-napthyridinyl-3-AICA) whose bulkier aromatic ring system led the RNA to crystallize under alternative conditions and in a different RNA conformation^40^. Residue C52 is observed adjacent to *m*-1-pyridinyl-AICA and must move to accommodate the bulkier ligand. Predictors were unable to predict this change in R1264 accurately, and no group achieved a local lDDT for C52 greater than 0.70 for R1264. However, a third of the models predicted the conformation C52 in R1263, which matched the template. 37% and 33% of models achieved lDDT for the RNA-ligand interface greater than 0.75. Predicting this type of conformational change, which is perhaps warranted given the size of the ligand, would greatly enable structure-guided design at the stage of ligand design and synthesis. This is particularly relevant owing to the difficulty in crystallizing bulkier ligands and the twinning apparent in these crystals.

### 2.5 SAM utilizing alkyltransferase ribozyme SAMURI (CASP: R1288, PDB: 9FN2) provided by Hsuan-Ai Chen and Claudia Höbartner

Most natural ribozymes catalyze the cleavage or formation of phosphodiester bonds, while synthetic ribozymes afford a broader spectrum of chemical reactions^41–43^. Recently, *in vitro* selected ribozymes that catalyze site-specific RNA modifications have been attracting increasing interest for their purported roles in an RNA world, and for their practical utility as tools in RNA biology. Specific examples of RNA-modifying ribozymes include the first methyltransferase ribozyme (MTR1)^44^, the engineered preQ1 riboswitch-ribozyme^45^, and two *S*-adenosylmethionine (SAM)-based ribozymes SMRZ1^46^ and SAMURI^47^. These ribozymes install methyl (and in some cases other alkyl) groups at defined positions within a target RNA.

Analyzing the structure of SAMURI is especially intriguing in comparison to natural SAM-binding riboswitches, which control gene expression in response to SAM or *S*-adenosylhomocysteine (SAH)^48^. While SAMURI readily transfers the methyl (or alkyl) group from SAM (or analogous selenomethionine cofactors) to N3 of a specific adenosine in the substrate RNA (**Figure 4A**), natural SAM riboswitches actively suppress RNA (self-)methylation. We solved the crystal structure of SAMURI in the post-catalytic state containing 3-methyladenosine (m^3^A) and SAH^49^. The structure revealed how SAMURI positions the reactants in an orientation poised for an SN2-like substitution reaction and, thus, adds a new example to the versatile structures RNAs can adopt to achieve distinct functions.

**Figure 4.**
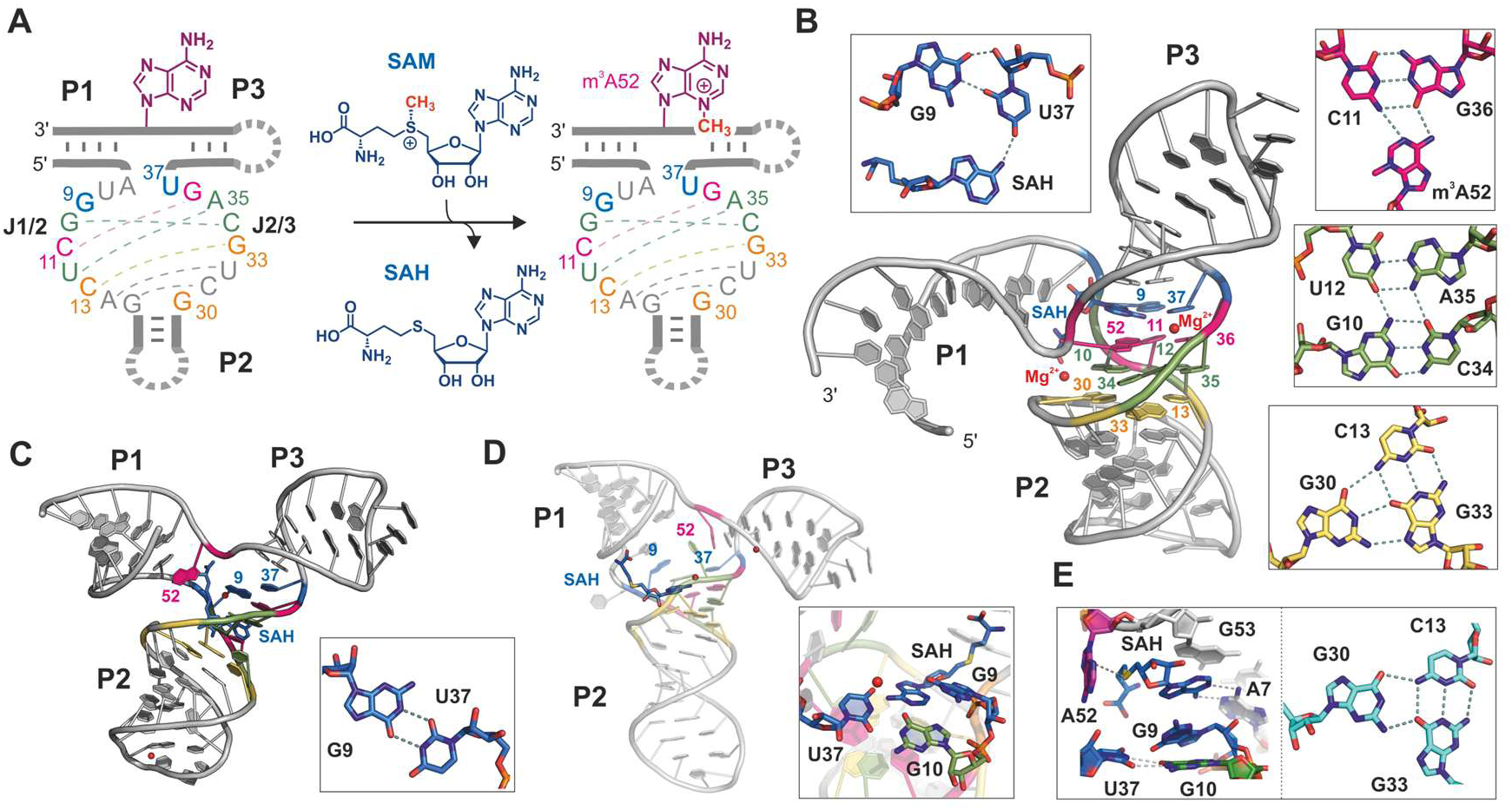
Prediction accuracy of SAMURI-SAM complex (CASP: R1288, PDB: 9FN2). (**A**) Secondary structure of SAMURI and the reaction scheme. Three A-form helices (P1-P3) are shown in bold lines and the catalytic core (J1/2 and J2/3) in colored letters. Colored dashed lines represent the Watson-Crick base pairs according to the crystal structure. (**B**) Crystal structure of SAMURI-SAM complex (PDB: 9FN2). Recognition patterns of each layer are shown in the boxes: cofactor layer in blue, reaction layer in magenta, stabilization layer in green and bottom layer in yellow. (**C**) The model R1288LG338_4 successfully predicted the coaxial folding of P2 and P3. A partially correct base pairing interaction in the cofactor layer is shown in the box. (**D**) The overall structure R1288LG262_4 and its SAH binding mode. (**E**) The predicted SAH binding mode (left box) and bottom layer (right box) from the model R1288LG055_1.

SAMURI adopts a three-way helix junction (3HJ) structure (**Figure 4A**), with the two substrate binding arms, P1 and P3, and the internal stem, P2, forming three canonical A-form helices. The catalytic core features a four-layered architecture composed of the cofactor layer, the reaction layer, the stabilization layer and the bottom layer (**Figure 4B**). A base triple in the bottom layer extends the π-π stacking of the coaxial P2 to P3 helices, which assists cofactor binding. A characteristic feature is the kink between P1 and P3 that docks the target adenosine (A52) in the reaction layer by H-bonding of its Watson-Crick and Hoogsteen edges, while exposing the minor groove edge to facilitate the reaction at N3.

This collection of key structural features of SAMURI was not recapitulated in the predictions. None of the R1288 predicted models within RMSD<10 Å managed to model the active site and ligand interactions correctly. The local distance difference test (LDDT) scored 0-0.16 for all models, which is lower than the general standard of a low-quality model (LDDT>0.3). This result reflects the challenge to produce the active site and ligand binding mode of synthetic functional RNAs.

Since the LDDT values are too low, we inspected the models with root mean square deviation (RMSD) within 10 Å (best score ∼4 Å), with the ultimate goal of solving an X-ray structure by molecular replacement using a predicted model (inspired by a similar approach by CASP15 evaluators^4^). We defined a manual scoring scheme to scrutinize the structural details of the overall fold with three helices, the four layers, the kink, the SAH binding and magnesium ion locations. As expected, all models predicted the three paired regions (P1, P2 and P3) correctly as double helices, albeit distinct in shape and orientation. Laudably, the models R1288LG338_4 and R1288LG272_1 successfully predicted the coaxial stacking of P2 to P3 (**Figure 4C**), even though the catalytic core was miscalculated in both cases. Despite several canonical base pairs in the catalytic core (**Figure 4A**, dashed lines), it was quite difficult for the algorithms to find the correct partners. The best predictions fell in the bottom layer, where half of the models correctly inserted G30 in a base triplet. The models R1288LG055_1 and R1288LG262_1 captured this interaction similar to the crystal structure (**Figure 4E**). Several of the models found the base pairs U12-A35 in the stabilization layer and C11-G36 in the reaction layer correctly. However, the correct partners for the experimentally observed triplet and quartet interactions were not identified. The sharp turn for locating A52 accurately in the backbone was another major hurdle. A few models (R1288LG272_1, R1288LG091_1 and R1288LG338_4) showed a slight turn but too shallow to form the reaction triplet (**Figure 4C**), while other models located A52 in a smooth transition from P1 to P3 (**Figure 4D**).

The location and binding mode of SAH were challenging to predict. While most models placed SAH near to the catalytic core, neither the orientation nor the binding mode were correctly calculated to reflect the G9 in *syn* conformation paired with the sugar edge of U37, which positions U37 to anchor SAH through H-bonding (**Figure 4B**) and stacking between G51 and A52. Only the model R1288LG338_4 spotted a G9-U37 interaction but predicted as a classical wobble pair (**Figure 4C**). Several other models placed G10 in a wobble pair with U37 instead of the experimentally observed Watson-Crick pair with C34 (**Figure 4D,E**), or predicted a G10-A35 purine-purine base pair in place of the stabilization quartet. The model R1288LG262_4 placed the SAH in a splayed conformation, with the base tucked in the core and the methionine tail pointing toward the opening space in P1 direction (**Figure 4D**), while model R1288GL055_1 found a solution with SAH base-paired to A7 and the homocysteine tail close to A52 (**Figure 4E**). In the crystal structure, two magnesium ions were found near the kink and the nucleobase of SAH (**Figure 4B**). The location of magnesium ions was unsuccessful in all predictions because both kink and SAH were incorrectly placed.

The best models according to manual scoring are R1228LG338_4, R1228LG272_1 and R1228LG091_1, although none reached more than 4 out of 10 points. Surprisingly, the rankings based on either LDDT or RMSD do not reflect the result of our assessment. Therefore, the top three candidates of each evaluation method (LDDT, RMSD and manual evaluation) were chosen to perform molecular replacement in Phenix^50^. Unfortunately, but not unexpectedly, none of these models generated a correct solution. The best translation function Z-score was 7, achieved by R1288LG338_4, which ranked high in our evaluation but was modest by LDDT and RMSD.

To summarize, the prediction of SAMURI structure in the product state with bound SAH was not successful, as reflected by the low LDDT score and medium to high RMSD values. Several models predicted the bottom triplet correctly and spotted two additional core base pairs accurately. However, none of the predicted models brought A52 and the ligand into an orientation that would infer the experimentally determined alkylation site. In general, structure predictions are still very challenging for synthetic RNAs that do not have any natural references and for which no analogous structures have previously been reported. Thus, more experimental structures are still needed to enrich the training repertoire.

### 2.6 Dopamine-bound DNA aptamer RKEC1 (CASP: D1273, PDB: 9HIO) provided by Eric Largy, Philip E. Johnson, and Cameron D. Mackereth

Aptamers are oligonucleotides that have been selected to bind a desired ligand, and many have been created to recognize small molecules for use in biosensors. Despite the increasingly popular development of small molecule-binding DNA aptamers, very little atomic information has been acquired on their ligand-aptamer complexes. This lack of molecular details thus limits our knowledge of binding mechanisms and hinders aptamer optimization. Related to CASP target D1273, a dopamine-binding aptamer was developed in the context of creating field-effect transistor-based sensors that rely upon ligand-induced DNA conformational changes^51^. Truncation of the duplex formed by the 5’ and 3’ ends did not significantly perturb binding characteristics^52,53^, and to create RKEC1 we have removed an additional three nucleotides at the 5’ end to further reduce the aptamer size and improve affinity.

Using NMR spectroscopy methods, we found that the dopamine-bound RKEC1 aptamer consists of a highly atypical DNA fold (**Table 1**; **Figure 5A,B**). The most notable feature of the DNA architecture is the absence of canonical duplex regions. In total, there are only two Watson-Crick base pairs (C14:G19 and C16:G23, numbered as in the CASP target sequence), and these are not adjacent in the structure. Despite the lack of canonical base pairs, there remain 11 additional base pairs formed by RKEC1 (solid lines in **Figure 5A**). These fall into categories more often present in RNA structures such as reverse Watson-Crick (T2:A25, T9:A17 and C10:G18), reverse wobble (T1:G24), and several homomeric pairs (G3:G26, A5:A25, G6:G24, G7:G23, G7:G24, G11:G19 and G11:G21). There are also six additional base pairs connected by a single hydrogen bond (dotted lines in **Figure 5A**). A final complexity in the structure arises from numerous places in which bases participate in two nucleotide planes, such that there are few instances of true base triples even though three DNA strands form several of the structural elements.

**Figure 5.**
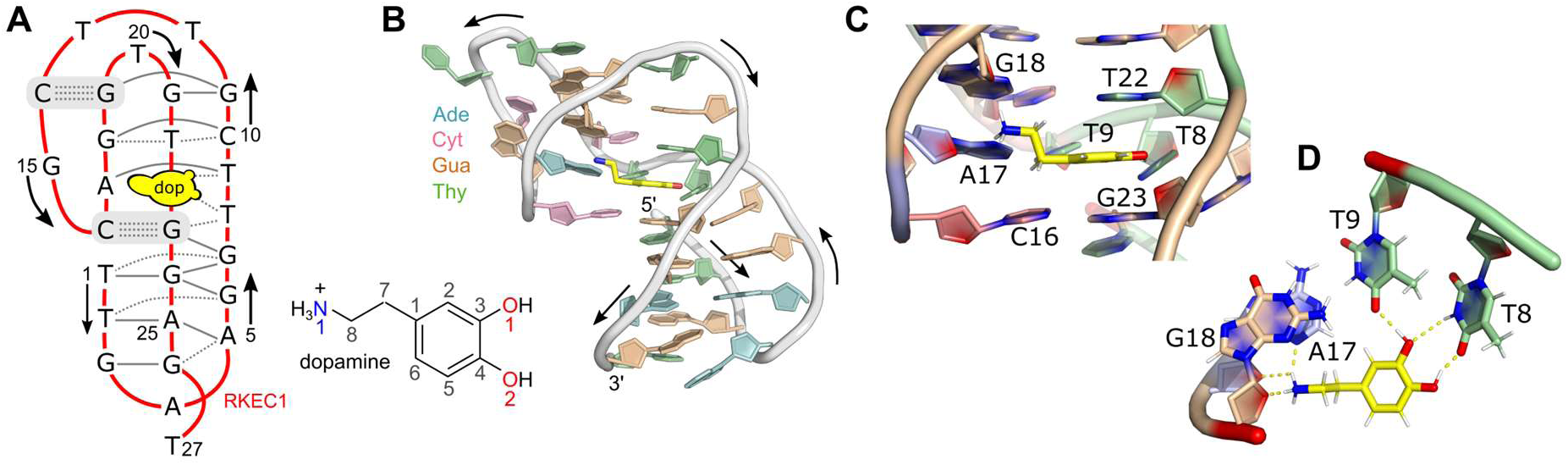
Molecular details of dopamine-bound RKEC1 aptamer (CASP: D1273, PDB: 9HIO). (**A**) Schematic of the dopamine-bound RKEC1 aptamer. The only two Watson-Crick base pairs are highlighted in gray. Additional base pairs with at least two hydrogen-bonds are shown as solid lines between nucleotides. Dotted lines indicate bases joined by a single hydrogen bond. (**B**) Representative cartoon model for the structure closest to the ensemble average of 15 calculated structures. (**C,D**) Illustration of the interactions the aptamer makes with dopamine due to stacking (**C**) and via hydrogen bonds (**D**).

Given the unusual nature of the RKEC1 DNA fold, it presents a difficult target. Predictions for the DNA in the complex included 107 models submitted to CASP from 22 different groups, of which 6 were obtained from servers. Despite the small size of the DNA at 27 nucleotides, none of the predicted models had an RMSD less than 10 Å, with model RMSD ranging from 10.16 Å to 27.79 Å. There are some broadly correct elements in the predictions, such as triple helical segments in some models and instances of bases that are shared between planes. When inspected in detail, however, the strand polarities differ from the experimental structure and bring together different bases. We had expected the atypical base-pairing to be challenging, but approximately 70 % of the 1254 base pairs in all models are formed by non-canonical base pair arrangements (i.e. not Watson-Crick). It is therefore possible that problems related to accurate secondary structure prediction were more inhibitory than the diversity of base pairing in RKEC1.

As there was dopamine present in the complex, there were also ligand predictions submitted. In this case, there were 8 groups that provided a total of 34 models. In the experimental structure, the dopamine ligand is positioned such that it stacks between bases T22 and G23 (**Figure 5C**). Hydrogen bonds connect the dopamine hydroxyls to T8 and T9, and the amine group by sugar and base atoms from A17 and G18 near the phosphate backbone (**Figure 5D**). In the predictions, the overall difficulty in predicting the DNA fold made evaluation of the ligand position difficult to assess using only statistics. By looking for similar types of contacts in the predictions, we noted some success in modeling the stacking interaction between dopamine and bases, but poor prediction of any hydrogen-bonding. For example, the five models from LG055 capture the correct idea of dopamine stacking between bases (but now placed between A4 and G24) and positions the dopamine amino group close to the phosphate backbone (between G3 and A4). In LG262 the three models also show a stacked dopamine between two bases (C10 and G21) but no other significant contacts. There are also several models with dopamine only stacked on one side (one model in LG262, four models in LG338, and four models in LG294) and interestingly dopamine is also stacked on one side even with single-stranded DNA (LG464).

Given the unusual DNA fold in the dopamine-RKEC1 complex, we anticipated that CASP target D1273 would be a challenging case. In terms of DNA structures in the PDB, there is an overrepresentation of duplex structures with the only other main contribution being depositions of G-quadruplexes. To date, there are only a small number of structures with higher DNA complexity, notably within some of the previously determined DNA aptamer structures. This deficiency in the availability of diverse folds likely plays a role in the lack of predictions with significant similarity to the experimental structure. It is also the case that for D1273 the dopamine ligand serves an integral part of the DNA fold: stacking between two bases and participating in hydrogen bonds. Prediction of just the DNA might therefore have also been difficult in the absence of the ligand.

### 2.7 *Oceanobacillus iheyensis* Group IIC intron (CASP: R1241) provided by Shekhar Jadhav, Michela Nigro, and Marco Marcia

Group II introns are bacterial and organellar self-splicing ribozymes that constitute the evolutionary ancestors of the eukaryotic spliceosomal small nuclear RNAs. Group II introns self-excise themselves from precursor mRNA transcripts in a two-step reaction that is identical in terms of molecular mechanism and chemistry to the reaction catalysed by the spliceosome^54^. In the first step of splicing, group II introns cleave their 5’-exon either through a hydrolytic or a transesterification mechanism, and in the second step of splicing the excised 5’-exon ligates to the 3’-exon producing a mature mRNA and an exon-free intron^55,56^. The free intron is still enzymatically active and can retrotranspose into genomic DNA target sites, with important biotechnological and medical implications in gene editing. To execute such complex reactions, group II introns assemble their six structural domains (D1 to D6, **Figure 6A**) into a well-ordered tertiary structure that harbours an RNA-based active site adjuvated by a metal-ion cluster cofactor in its core (**Figure 6B**)^54^. This active site can be specifically and selectively targeted by small molecules with potential applications as antifungals^57^. Resolving the molecular architecture of group II introns accurately at near-atomic resolution and elucidating their precise mechanism of action is therefore crucial to understand the fundamental reaction of splicing, to support the design of antifungal drugs, and to guide the engineering of novel RNA nanomachines for gene editing.

**Figure 6.**
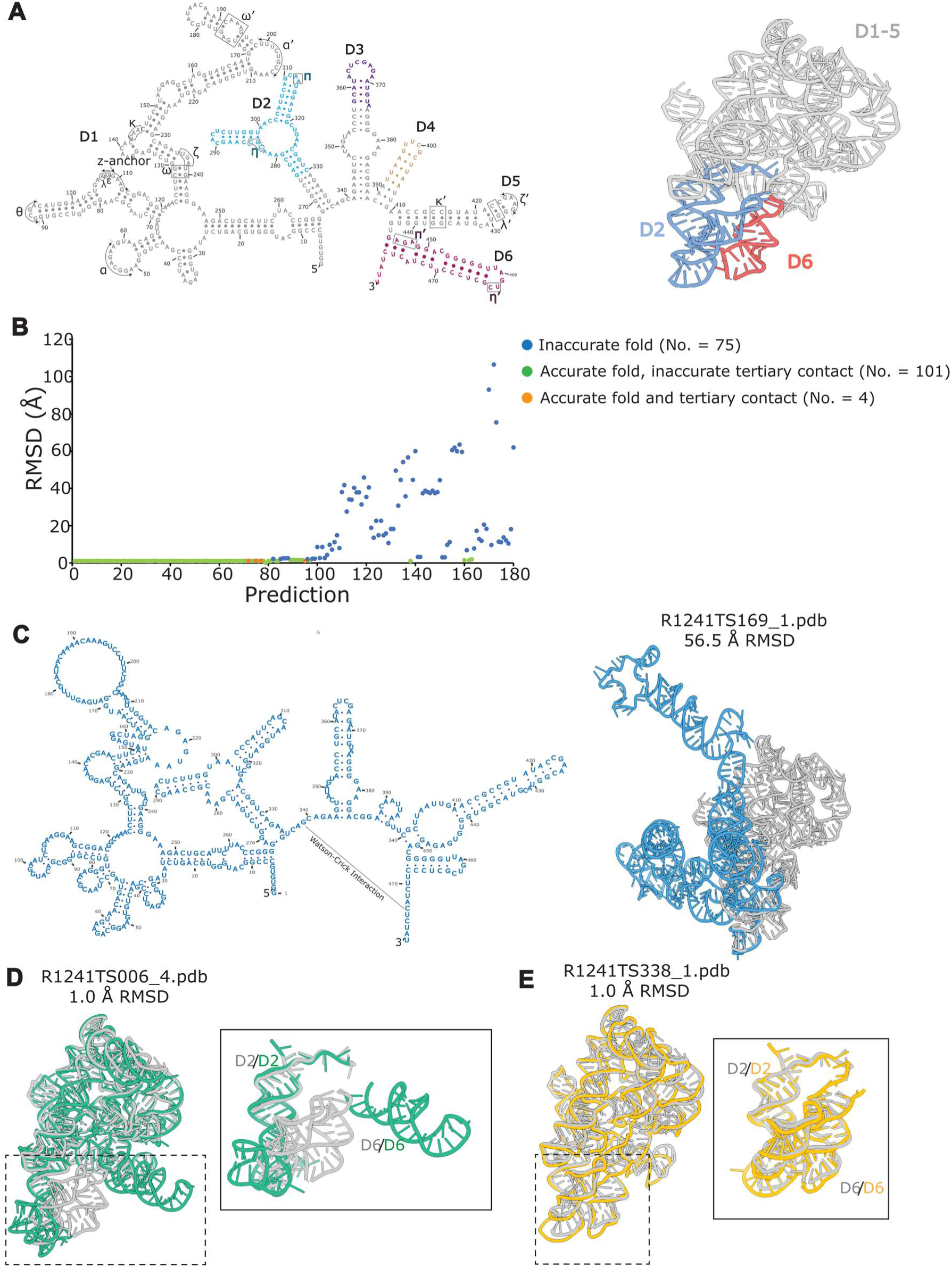
***O. iheyensis* group II intron structural analysis (CASP: R1241).** (**A**) Secondary and tertiary structures of *O. iheyensis* group IIC intron. D1-D6 indicate structural domains 1 to 6, respectively. The previously unresolved regions in D2 and D6 are coloured in blue and Indian red, respectively. The tertiary contacts are indicated with Greek letters. (**B**) All-atom RMSD distribution of the 180 structures predicted in CASP16 with respect to our experimental model. Blue points display the 75 inaccurate structures. Green points display the 101 structures that show accurate prediction of the D1-5 fold but inaccurate predictions of D2-D6 contacts. Orange points display structures that are accurate in overall fold and local D2-D6 tertiary contacts. (**C**) Secondary and tertiary structures of the R1241TS169_1 prediction, representative of the 75 inaccurate predictions. The tertiary structure of R1241TS169_1 (in blue) is displayed superposed over our experimental model (in grey). (**D**) Superposition of our experimental model (in grey) over the R1241TS006_4 prediction (green, representative of the 101 structures with inaccurate fold by inaccurate D2-D6 contacts). The inset displays a zoom into the D2-D6 interaction. (**E**) Superposition of our experimental model (in grey) over the R1241TS338_1 prediction (orange, representative of the 4 structures with accurate fold and D2-D6 contacts). The inset displays a zoom into the D2-D6 interaction. Images of RNA 3D structure and secondary structure map were generated in the ChimeraX and VARNA software respectively^59,60^.

So far, experimental structures are available for domains D1-D5 of the so-called group IIC intron from the bacterium *Oceanobacillus iheyensis*, which splices through the hydrolytic mechanism, and for D1-D6 of homologous group IIA and IIB introns from other bacteria or algal chloroplasts, which splice through the transesterification mechanism^54^. The structure of D6 from the *O. iheyensis* group IIC intron is instead still undetermined. As a result, it remains unclear how D6 – which harbors key functional elements such as a branch-point adenosine and the 3’-splice junction – interacts with the rest of the ribozyme in introns that splice through the hydrolytic mechanism. We determined the D1-D6 structure of the *O. iheyensis* group IIC intron by cryo-electron microscopy (cryo-EM) in the resolution range between 3.3 – 4.1 Å (3.8 Å average resolution), and provided its sequence as a target for the CASP16 competition.

36 groups participated in the competition and presented 180 model structures of our target. Here, we are discussing the predicted models in comparison to our experimental cryo-EM structure with respect to the following parameters: (i) accuracy in the prediction of the secondary structure; (ii) accuracy in the prediction of the overall tertiary fold, as judged by the all-atom root mean square deviation (RMSD) between predicted and experimental models; and (iii) accuracy in the prediction of specific tertiary contacts, with a focus on the previously-undetermined contacts between D6 and the rest of the molecule (known from biochemical characterization as the η-η’ and π-π’ interactions between D6 and D2)^54^.

75 predictions display inaccurate tertiary structure models, with RMSD values of 2-120 Å with respect to the experimental model (**Figure 6C**). Most of these predictions crucially fail in correctly reproducing the experimental secondary structure of the target (**Figure 6D**). Alternatively, those predictions that reproduce the correct secondary structure fail in modelling the characteristic five-way junction of D1, which is the folding scaffold that dictates the orientation of all peripheral elements with respect to the core^58^.

The remaining 105 predictions display an accurate secondary structure and overall tertiary fold, with RMSD <2 Å with respect to the experimental structure (**Figure 6C**). However, 101 of these predictions fail to accurately visualize the novel D6 η-η’ and π-π’ interactions. Among these structures, 21 structures display no interaction at all between D6 and the rest of the ribozyme and 80 structures display tertiary interactions between D2 and D6 different from the experimentally-validated η-η’ and π-π’ (**Figure 6F-G**). A characteristic feature of the predictions that fail to reproduce the experimental D2-D6 interactions is that they model inaccurately the characteristic 3-way junction of D2 (**Figure 6H-J**).

Remarkably, 4 predictions display accurate models in terms of secondary and tertiary structure organisation and of D2-D6 interactions. Among these models, three were predicted by the Vfold group and one was predicted by the GeneSilico group. Vfold uses the Protein Data Bank (PDB) server-based structure templates to produce an initial *ab initio* model, which is then refined by all atom energy minimization^61^. Its success on our target can thus possibly be explained by the existence of 12 group IIA/B intron structures homologous to our target in the PDB (41-56% sequence identity) and displaying the D2-D6 interaction. Compared to Vfold, GeneSilico uses coarse-grained RNA representations and Monte Carlo sampling to explore the RNA conformational space and derive the most energetically-favourable model^62^. This approach seems to be the most unbiased and accurate on our target.

In summary, our analysis reveals that key factors of success in RNA structure modelling are the accurate predictions of RNA secondary structure motifs (stem loops), multi-way junctions and long-range tertiary contacts. A current limitation resides in the fact that prediction algorithms currently do not predict metal and ligand binding sites. This limitation is particularly severe for our target, which harbours a functional metal cluster in the active site and additionally requires tens of rigidly-bound ions for structural stability^63^. We suggest that future generations of RNA structure modelling algorithms include ion cofactors as intrinsic tertiary structure elements of the targets to improve RNA structure predictions^64^.

### 2.8 Rumen-Originating, Ornate, Large (ROOL) RNA hexamer (CASP: R1286, PDB: 9J6Y) and octamer (CASP: R1283v1-v3, PDB: 9ISV, 9J3R and 9J3T) provided by Liu Wang, Jiahao Xie, and Zhaoming Su

The Rumen-Originating, Ornate, Large (ROOL) RNA, one of the largest and most structurally complex non-coding RNAs, was first identified through comparative analysis of cow rumen metagenome data^65^. Another independent study revealed its presence in the megaplasmid of *Lactobacillus salivarius* (*Lsa*), where its abundance can exceed that of 16S ribosomal RNA under specific conditions^66^. ROOL RNA frequently localizes in prophage regions, purified phage particles, non-prophage regions, and tRNA loci^65^. The biological functions of ROOL RNAs remain unclear. Recent studies have resolved the cryo-EM structures of ROOL RNAs from *Enterococcus faecalis* (*Efa*) as 3.1 Å monomer (R1283v1), 4.7 Å tetramer (R1283v2) and 3.8 Å octamer (R1283v3), from *Lsa* as 3.1 Å hexamer (R1286), and from environmental bacteria (*env-120*) as 3.1 Å octamer (**Table 1**, **Figure 7A-B**)^67,68^. Another two ROOL RNA sequences from *Lsa* forming a hexamer (CASP: R1252) and environmental bacteria (*env-*209) forming octamers (CASP: R1253) with 2 alternative conformations have also been submitted for CASP16^68^. Except for the two *Lsa* sequences, the other sequences share less than 50% similarity. However, they all adopt a conserved architecture comprising 16 base-paired regions (denoted P1-P16, **Figure 7C**), five multiway junctions, and intricate interaction networks involving base-pairings, minor groove interactions and magnesium ion coordination.

**Figure 7.**
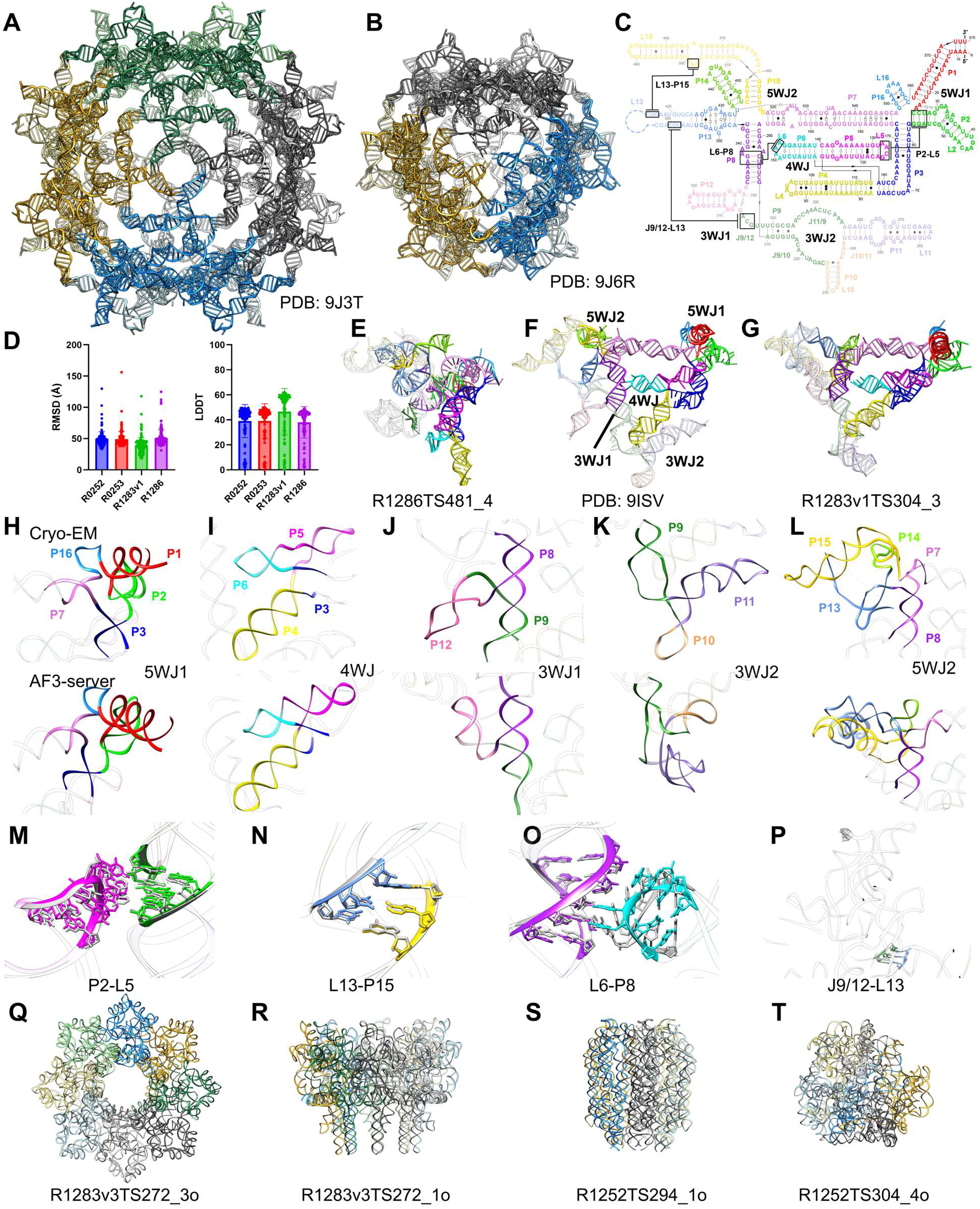
Evaluation of the ROOL RNA structure predictions (CASP: R1286 and R1283v1-v3, PDB: 9J6Y, 9ISV, 9J3R and 9J3T). (**A-B**) Cryo-EM structure of *Efa* ROOL octamer (R1283) and *Lsa* ROOL hexamer (R1286). (**C**) Secondary structure of *Efa* ROOL monomer, with flexible regions P9-P12, P13, and P15 in transparency. Intramolecular interactions are highlighted by lines and boxes. The unresolved region L13 is annotated with a dotted line. (**D**) Average RMSD and LDDT of ROOL monomers, of R0252, R0253, R1283v1 and R1286. (**E**) Representative predicted model of *Lsa* ROOL. (**F**) Cryo-EM structure of *Efa* ROOL monomer with regions resolved only in oligomeric states annotated in transparency. (**G**) Predicted model of *Efa* ROOL by AF3_server with consistent secondary structure. (**H-L**) Comparison of multiway junctions between cryo-EM and R1283v1TS304_3 predicted structures of (**H**) 5WJ1, (**I**) 4WJ, (**J**) 3WJ1, (**K**) 3WJ2 and (**L**) 5WJ2 in *Efa* ROOL monomer. (**M-P**) Superposition of intramolecular interactions between cryo-EMand R1283v1TS304_3 predicted structures of (**M**) P2-L5 TL/TLR, (**N**) L13-P15 PK, (**O**) L6-P8 minor-groove and (**P**) J9/12-L13 PK interactions in *Efa* ROOL monomer. (**Q-T**) Predicted homo-oligomers of ROOL RNAs forming (**Q**) ring-like structure (R1283v3TS272_3o), (**R**) bundle-like structure (R1283v3TS272_1o), (**S**) RNA origami-like structure (R1252TS294_1o) and (**T**) unusual asymmetric structure (R1252TS304_4o).

Predictions for the four ROOL RNA monomer structures are generally inaccurate, as evidenced by average RMSDs worse than 40 Å and average LDDTs below 50 (**Figure 7D**). Although comparative genomics analysis has shown a highly conserved secondary structure^65^, the majority of predictions has not incorporated sequence conservation and covariation into their predicted 3D structures, thus not further discussed (an example in **Figure 7E**). The high flexibility of ROOL RNA monomer structure could be one reason for the poor prediction results, as demonstrated by the missing density in the *Efa* ROOL monomer structure that is only resolved in oligomeric states (**Figure 7F**). While none of the prediction results resemble the experimental structures in R0252, R0253 and R1286, an interesting result appears in R1283v1 monomer prediction, in which the AlphaFold 3 model R1283v1TS304_3 submitted by the AF3-server exhibits a highly accurate secondary structure and a reasonable overall fold compared to R1283v1 (GDT-TS = 0.15, TM-score = 0.365, LDDT = 0.619) (**Figure 7G**).

The predicted model consists of P1-P16 and contains all five multiway junctions in consistency to the conserved secondary structure and cryo-EM structure (**Figure 7H-L**). The first five-way junction (5WJ1) consisting of P1-P3, P7 and P16, and the four-way junction (4WJ) consisting of P3-P6 closely resemble the cryo-EM structure (**Figure 7H-I**), whereas P12 in the first three-way junction (3WJ1) containing P8, P9 and P12, is translocated away compared to the cryo-EM structure (**Figure 7J**). The predicted 3WJ2 and 5WJ2 do not agree with the cryo-EM structure, in which 3WJ2 contains swapped P10 and P11 (**Figure 7K**), whereas the 5WJ2 contains an incorrect P13 and swapped P13 and P15 (**Figure 7L**). Intriguingly, the AF3-server result accurately predicts two intramolecular tertiary interactions, P2-L5 tetraloop/tetraloop receptor and L13-P15 pseudoknot (PK) (**Figure 7M-N**), and closely resembles the L6-P8 minor-groove interaction present in the cryo-EM structure (**Figure 7O**). The last intramolecular PK L13-P15 is not predicted (**Figure 7P**). This reasonable prediction of such a complex RNA structure is most likely achieved by taking the sequence conservation and covariation into account during structure prediction. All other predictions of *Efa* and other ROOL RNAs did not yield similar structures. Perhaps, the AF3-server prediction relied on models that were sampled using automatically generated multiple sequence alignments in which more sequences were included for R1283 resulting in a more complete covariation analysis than other ROOL sequencesCASP16 NA assessment paper.

Homo-oligomeric protein structures have been extensively characterized. In contrast, there are far fewer examples of oligomeric RNA structures, and RNA homodimers are most commonly reported^69–71^. In fact, there are only four homo-multimer structures from two RNA sequences in the PDB consisting of more than two protomers, including the φ29 prohead RNA forming either four- or five-membered ring architectures, and the Chili RNA aptamer tetramer^72–74^. This scarcity of RNA oligomer structural information has compromised the prediction accuracy of RNA oligomers in CASP16. We speculate that most oligomer predictions rely on existing RNA and protein homo-oligomer templates in the ROOL hexamer or octamer prediction tasks (R1252o, R1253v1, R1253v2 and R1283v3o). These predictions include ring-like structures resembling the φ29 prohead RNA or pore-forming protein oligomers, bundle-like structures that look close to membrane protein bundles and some synthetic RNA origami (**Figure 7Q-S**). While most predictions for ROOL oligomers are more or less symmetrical, we also notice unusual cases of asymmetric oligomeric assembly (**Figure 7T**).

### 2.9 Ornate Large Extremophilic (OLE) RNA Dimer (CASP: R1285o, PDB: 9MCW) provided by Rachael C. Kretsch, Yuan Wu, Rhiju Das, Wah Chiu

The Ornate Large Extremophilic (OLE) RNA family was identified as a large, non-coding, highly structured RNA family in gram-positive bacteria in extreme environments by covariation analysis^75^. Prior to CASP16, OLE RNA was known to form a RNA-protein complex which localizes to the cell membrane^76^ and was implicated in many environmental sensing pathways^77^. However, the biochemical mechanism of action remains unknown^78^.

The 5’ domain of OLE RNA from *Halalkalibacterium halodurans* forms a well-ordered RNA-only dimer which resolves to 2.9 Å using single particle cryogenic electron microscopy^68^ (**Table 1**, **Figure 8A-B**). Each monomer has five helices (P4-P8) which lay parallel to each other such that the loops of the helices point into the dimer interface. These loops form two dimer interfaces labeled B1 and B2. The final helix (P9) crosses this stack perpendicularly, eventually pointing into the dimer interface and forming the final dimer interface, B3. Two independently solved cryo-EM structures of the OLE RNA dimer agree closely, supporting the use of this molecule as a target for CASP16 prediction^67,68^. The stoichiometry was given as a dimer to predictors.

**Figure 8.**
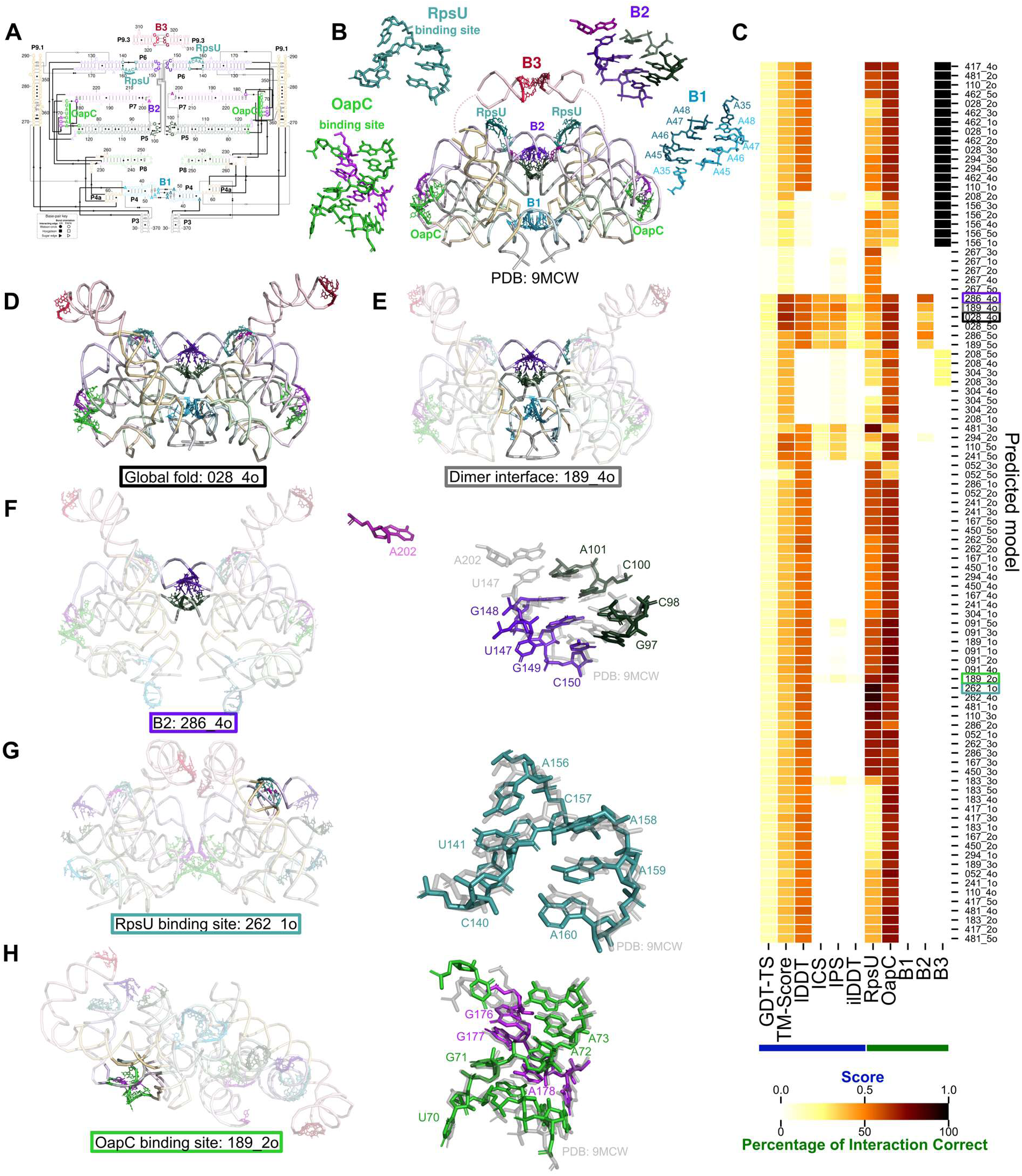
Predictions of the OLE RNA dimer (CASP: R1285o, PDB: 9MCW). (**A**) The secondary structure of the OLE RNA dimer is displayed with one monomer on the left, another on the right, and the dimer interface in the center. (**B**) The cryo-EM structure of the OLE RNA dimer, PDB: 9MCW, is displayed in the center in ribbon format. The two protein binding sites and three dimer interfaces are labeled and the nucleotides involved are displayed. Zoom-in of four of these areas of interest surround the model. (**C**) All submitted models for R1285o are cluster by performance based on various metrics: global fold metrics (GDT-TS, TM-score, lDDT), interface metrics (ICS, IPS, ilDDT), and percent of native interactions of interest that were recovered. Interactions in the cryo-EM structure and the predicted models were labeled by rna_motif^80^. For the intermolecular interactions, only inter-chain interactions were counted. Models of interest are boxes and their structures are displayed in (**D-H**).

No homologous structures are known, posing a challenge for prediction. However, there is a large multiple sequence alignment and biochemical information in the literature. According to the CASP16 metrics, 028_4o had the most accurate fold (GDT-TS = 23.2, TM-score = 0.682, lDDT = 0.572) while 189_4o had the most accurate dimer interface (ICS = 0.444, IPS = 0.532, ilDDT = 0.297) (**Figure 8C-E**). Although these models predicted the global fold, the predictions are inaccurate in important structural and functional regions: dimer interfaces and protein binding domains.

At most one of the three dimer interfaces was predicted correctly in each model and the majority of models predicted no interface correctly. 20 models predicted B3 correctly which is the simplest interface made of a kissing-loop with four G-C base pairs (**Figure 8C**, this region was low resolution and hence not assessed in the official CASP16 assessment). Only 6 models predicted a B2-like interface, none of which predicted accurate base interactions (**Figure 8C**). B2 is a set of base-pairing and stacking interactions between the loops of P5 and P7 of one chain and the loop of P6 of the other chain. Model 286_4o modeled the interaction between the loops of P5 and P6 as the only intermolecular interactions, but it did not predict the additional involvement of the loop of P7 (**Figure 8F**). B1, a set of four non-canonical A-A base-pairs between the loops of P4 of both chains, proved the most challenging interface for predictors. Although some models predicted these loops in close proximity (**Figure 8D-E**), none predicted the base pairing between the loop (**Figure 8C**). The dimer interface could play an important role in scaffolding the OLE RNA-protein complex or as a switch in the OLE RNPs response to external stimuli. Therefore, it is important to improve the accuracy of RNA-RNA interface predictions to foster a better understanding of these biological systems.

Within each monomer, there are two protein binding sites that appear preorganized to expose the known protein binding sequences but, in both sites the RNA secondary structure differs from that hypothesized in the literature^79^. First, RpsU is known to bind a CAA motif in P6 that was previously thought to be partially precluded in a stem. Contrary to this, the cryo-EM structure shows that this motif is solvent exposed, potentially allowing for protein docking without RNA structural rearrangement. The majority (61%) of models incorrectly predict at least two of these three nucleotides are paired, although a few models did predict this site relatively accurately (**Figure 8C,G**). Finally, OapC was known to bind a kink turn formed immediately upstream P5. It was previously hypothesised that this region base-paired with the region immediately downstream of P5, however, the cryo-EM structure shows that three G-A non canonical base-pairs were formed with the region between P6 and P7 instead. 75% of the models predicted the correct three G-A non canonical base-pairs with many additionally obtaining excellent accuracy for the kink turn (**Figure 8C,H**). This observation suggests that predictors were able to use the sequence or evolutionary data to deduce the residues involved in the kink turn.

The analysis of OLE RNA predictions was encouraging, as the global fold was successfully predicted despite the absence of homologous RNA; however, critical interactions were still inaccurately predicted. In particular, predictors were not able to predict the dimer interface, modeling at most one of the three interfaces. In fact, many of the models predicted only one or no RNA-RNA interaction between the chains, underpredicting the complexity of the interface. It was also evident that predictors could only successfully predict the simplest type of interface, canonical base-pairing, with few or no correct interactions predicted for the more complex interfaces. Hence, structure predictors should aim to improve RNA-RNA interface prediction which would have implications for understanding the broader OLE family of RNAs and aid in the identification of other RNA families that form similarly complex interfaces.

### 2.10 RNA origami dimer structure and protein-binding aptamer with 2’F pyrimidine modifications (CASP: R1281o and M1282) provided by Emil L. Kristoffersen, Nikolaj H. Zwergius and Ebbe S. Andersen

Production of nucleic acids using chemically modified building blocks has proven essential for medicine and biotechnology. Indeed, modified nucleic acids, including natural as well as unnatural modifications of the canonical nucleotides^81^, are used in mRNA vaccines^82^, RNAi^83^ and gene editing^84^. For these reasons, it is becoming increasingly important to be able to model and predict the structure and effect of modified nucleic acids.

2’ Fluoro (F) modified RNA, is a widely used type of modification that greatly improves polymer properties. However, while the highly electronegative F modification increases polymer stability as well as base pairing and stacking stability, it alters the interactions and the structures that can be formed^85^. A blend of 2’ F pyrimidines (FY) and natural purines (termed 2’ FY RNA) is therefore often used to make a very RNA-like polymer, with greatly increased serum stability and efficient production strategies using *in vitro* transcription with a mutant RNA polymerase^86^. However even 2’ FY RNA alters folding.

We have produced and investigated 2’ FY RNA structures by *in vitro* transcription and cryogenic electron microscopy (cryo-EM). For CASP16 we uploaded the sequences of 1) an 2’ FY RNA origami design (R1281) which were noted to have A2-stoichiometry and 2) an 2’ FY RNA protein-binding aptamer and its binding partner (the RBD domain of the SARS-CoV2 spike protein) (M1282). Both targets had been investigated with cryo-EM revealing distinct 3D structures when produced in FY RNA compared to RNA. Our structural analysis allowed determination of the strand-path and general folding for the origami (GSFSC resolution ∼10 Å) and precise structural determination of the aptamer-protein complex (GSFSC resolution ∼4 Å).

Importantly, the CASP target submission platform uses FASTA format and does not allow specification of modified nucleotides or amino acids. Therefore, we noted in “additional information” that the RNA sequence had 2’ F modified pyrimidines. But in the end, all predictions were performed with regular RNA.

#### 2’ FY RNA origami dimer (R1281o, stoichiometry A2)

We produced a 6-helix origami structure of 2’ FY RNA, which has previously been investigated as RNA^87^. It was originally designed to form a bundle-like structure, with two RNA helices closing the bundle by forming a “clasp” held together by kissing loop interactions. Interestingly the RNA origami had a slow maturation process, going from a metastable young conformation to a later forming mature conformation^87^. The RNA version of the origami was predicted in CASP15 and is now on the protein data bank (PDB: 7PTK (young) and 7PTL (mature)). Surprisingly, we found that the 2’ FY RNA version of the origami dimerized. Despite relatively poor resolution (∼10 Å) we were able to confidently trace the backbone and build a general structure of the dimeric origami (**Figure 9A**). The model shows that the molecule rearranges so that two helices in each monomer form two intermolecular kissing loops instead of the intramolecular kissing loop observed in the RNA version.

**Figure 9.**
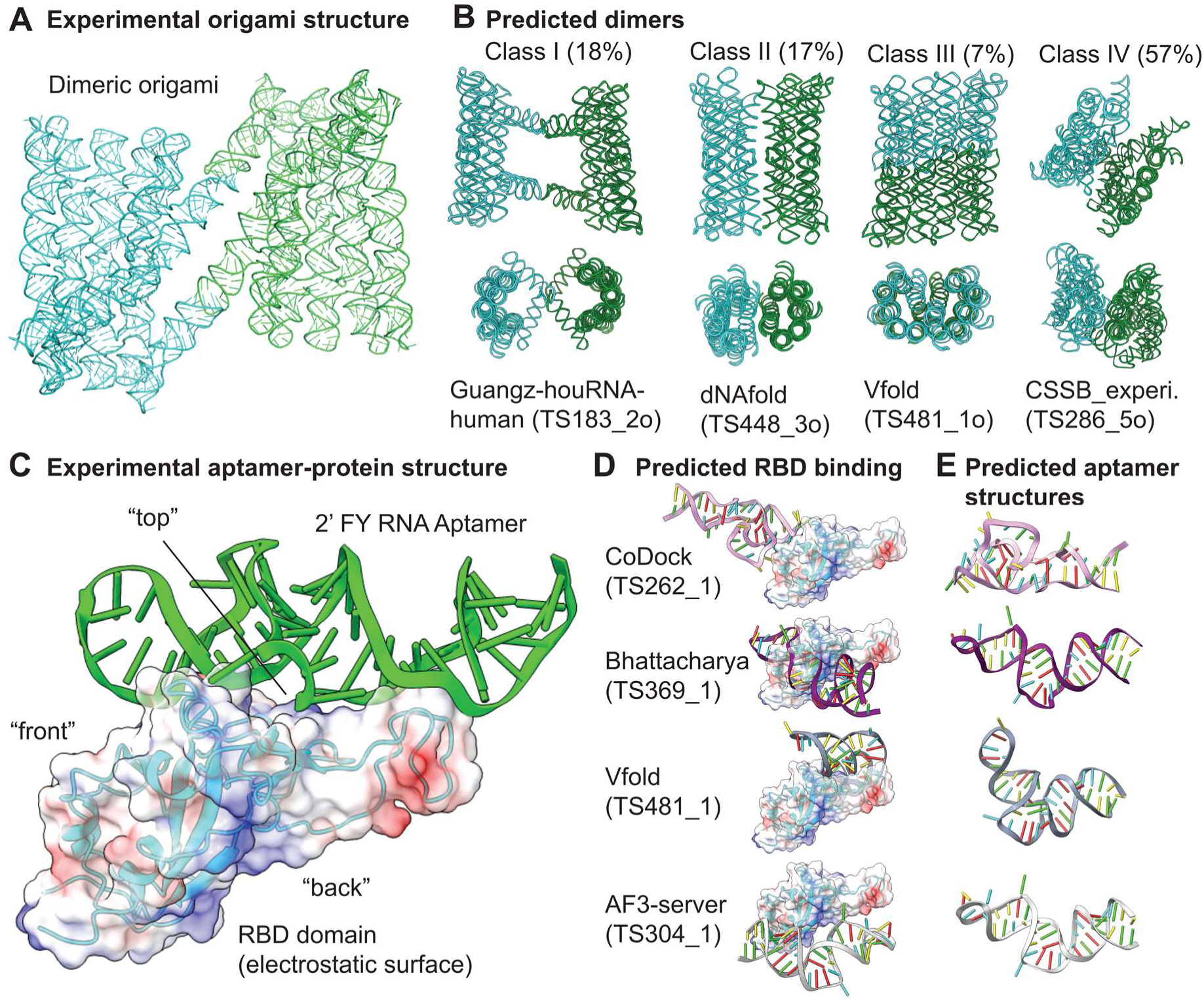
Prediction of 2’ FY RNA dimeric origami and 2’ FY anti-spike aptamer (CASP: R1281o and M1282). (**A**) Experimental determined structure of the dimerized 2’ FY RNA 6 helix bundle with a clasp origami. (**B**) Examples of predictions classified into class I to IV based on manual assessment. (**C**) Experimental determined structure of the 2’ FY aptamer (green) binding to the RBD domain (blue cartoon with transparent surface representation of electrostatics). (**D**) Examples of predictions where the aptamer binds different regions of the RBD domain (aligned according to the RBD domain). (**E**) The aptamers alone.

CASP16 provided 95 predictions of this homo dimer. We accessed all predictions and classified them into four classes based on manual assessment (I-IV) (**Figure 9B**). Class I (18%) consisted of models looking like the published RNA origami bundle (resembling the young structure) but where the clasp formed an intermolecular kissing loop. At first glance Class I looked promising, however a closer look revealed that the intermolecular kissing loops were not the correct ones, as observed in the experimental structure. Class II (17%) consisted of predictions where two bundle-like structures (resembling the mature RNA bundle) were positioned next to each other in different ways, without forming any base pairing interactions. Class III (7%) consisted of a split-open origami that dimerizes with all its internal kissing loops. Such a structure might be interesting to design in the future but did not resemble the experimental structure. Class IV was the most populated class (57 %) representing somewhat collapsed structures looking very different from the experimental origami structures.

In summary, none of the predictions captured the rearrangement that happened in the 2’ FY RNA origami causing it to dimerize. Importantly, the predictions were submitted as unmodified RNA. However, only the 2’ FY RNA version of the bundle dimerizes. Indeed the ability to specify modifications and use them in the predictions might be beneficial.

#### 2’ FY RNA aptamer / protein complex (M1282)

A 2’ FY RNA aptamer binding to the SARS-CoV 2 spike protein had previously been reported^88^, but its precise binding and structure was unresolved. We investigated its structure with cryo-EM, revealing that the aptamer binds to the Receptor Binding Domain (RBD) of the spike protein (**Figure 9C**). Importantly, an RNA version of the aptamer does not bind the spike protein at all.

CASP16 provided 31 predictions of the aptamer / RBD binding. We first accessed all predictions in terms of binding location (**Figure 9D**) and then the aptamer structure (**Figure 9E).** 56 % of the predicted aptamers bound to the positively charged “back” of the RBD domain (see electronegative colored surface **Figure 9D**, lower right). 8 % bound the “front” of the RBD domain (as seen in **Figure 9D** top left). 36 % of the aptamers were predicted to bind to the the “top” of the RBD domain (**Figure 9D** lower left), which is the region where the aptamer binds according to the experimental data. Despite binding to the correct regions of the RBD, these predictions did not identify the specific interactions between the nucleic acid molecule and the protein. Next we aligned all the aptamers (discarding the RBD domain from the models). This alignment revealed that most predictions identified the correct base pairing pattern. But none of them predicted the precise tertiary interactions (**Figure 9E**).

In summary, the base pairing patterns of the aptamer were often predicted correctly, however important tertiary interactions were overlooked. Due to the missed tertiary interactions, and / or other factors, the correct binding was, despite sometimes locating to the correct area of the RBD, not predicted very well. Again the predictions were done for RNA (not 2’ FY RNA). It is important for future prediction of chemically modified biomolecules to be able to include these. However there are very many different modifications, and it might be a difficult task to achieve. 2’ F RNA might be a good place to start.

### 2.11 Cryo-EM structure of SPβ LSI-RDF synaptic complex from the Bacillus phage SPβ (CASP: M1239v1 and M1239v2, PDB: 9DXD (pre-rotation state)). Provided by Heewhan Shin and Phoebe A. Rice

Large serine integrase (LSI) and recombination directionality factor (RDF) pairs are encoded by many temperate phages. Upon infection, the LSI catalyzes the unidirectional integration of the phage DNA into that of its bacterial host, where it can passively “hitchhike” for many generations. When the phage needs to escape that host and find a new one, it expresses the cognate RDF protein that binds to the integrase, altering its preferred reaction direction and triggering excision of phage DNA out of the host genome.

LSI/RDF pairs are of interest as fundamentally intriguing molecular machines and as biotechnology tools. In addition to high site-specificity and unidirectionality in DNA recombination, LSI/RDF systems display several advantages over other commonly used genome editing tools^89,90^. For example, LSIs require much shorter DNA binding sites than integrases from the tyrosine recombinase family. Compared to CRISPR-RNA guided systems, not only are the LSIs intrinsically designed to operate multiplexed insertion/deletion/inversion of kilobase-sized DNA payloads, but also no DNA repair is needed after recombination reaction because all broken bonds are relegated.

LSIs have a modular structure with an N-terminal catalytic domain followed by two DNA binding domains, with a coiled coil (CC) inserted into the 2^nd^ DNA binding domain (DBD2) via flexible hinges. They bind as dimers to their cognate DNA sites, but recombination requires formation of a tetrameric complex that synapses the two substrate DNAs. The substrate DNA sites for integration are termed *attB* (∼40 bp; from the bacterial host genome) and *attP* (∼50 bp; from phage DNA)^89,91,92^. The products of integration are two sites (“*attL*” and “*attR*”) that each contain one half of *attB* and one half of *attP*. *AttL* and *attR* are the substrates for the excision reaction, which regenerates *attB* and *attP*. The critical difference among these sites, which dictates whether inactive dimers or active tetramers can be formed, is the placement of the DBD2-binding motif, which is 4-5 bp further from the center in *attP* sites vs. *attB* sites. While the catalytic domain is central to both dimers and tetramers, additional interactions between the tips the CCs, which move with DBD2, form “handshake” interactions that stabilize active tetramers before recombination but also lock product dimers to prevent reverse reactions.

We were particularly focused on understanding the mechanistic basis behind how LSIs and their cognate RDFs control the directionality of DNA recombination. Previous biochemical and structural studies described the importance of the CC motif and the functional difference between *attB* and *attP* sites^93^, yet there were no available structures of intact LSI-DNA synaptic complexes, nor were there models showing how the RDF controls the CC’s trajectory. Our recent series of structures fill these gaps^94^.

We thought that the synaptic complexes would be interesting CASP targets due to their conformational complexity (**Figure 10A left**). First, while the four LSI molecules are chemically identical, their relative conformational arrangements differ based on which DNA half sites (attB- vs attP-derived) they are bound to, creating structural asymmetry; second, this asymmetry positions the tips of the CC-motifs to make tetramer-stabilizing in inter-dimer interactions (only between subunits bound to different types of half-sites); third, we captured the complex in the recombination intermediate state where each catalytic Ser22 residue is covalently linked to the 5’ end of the phosphate at the cleavage site. Finally, the catalytic domains can switch between inactive dimer and active tetramer conformations. Here we analyze the CASP results specifically for the complex of phage SPβ integrase, with its cognate RDF, synapsing two copies of *attL* - essentially, the complex poised to undertake the excision reaction.

**Figure 10.**
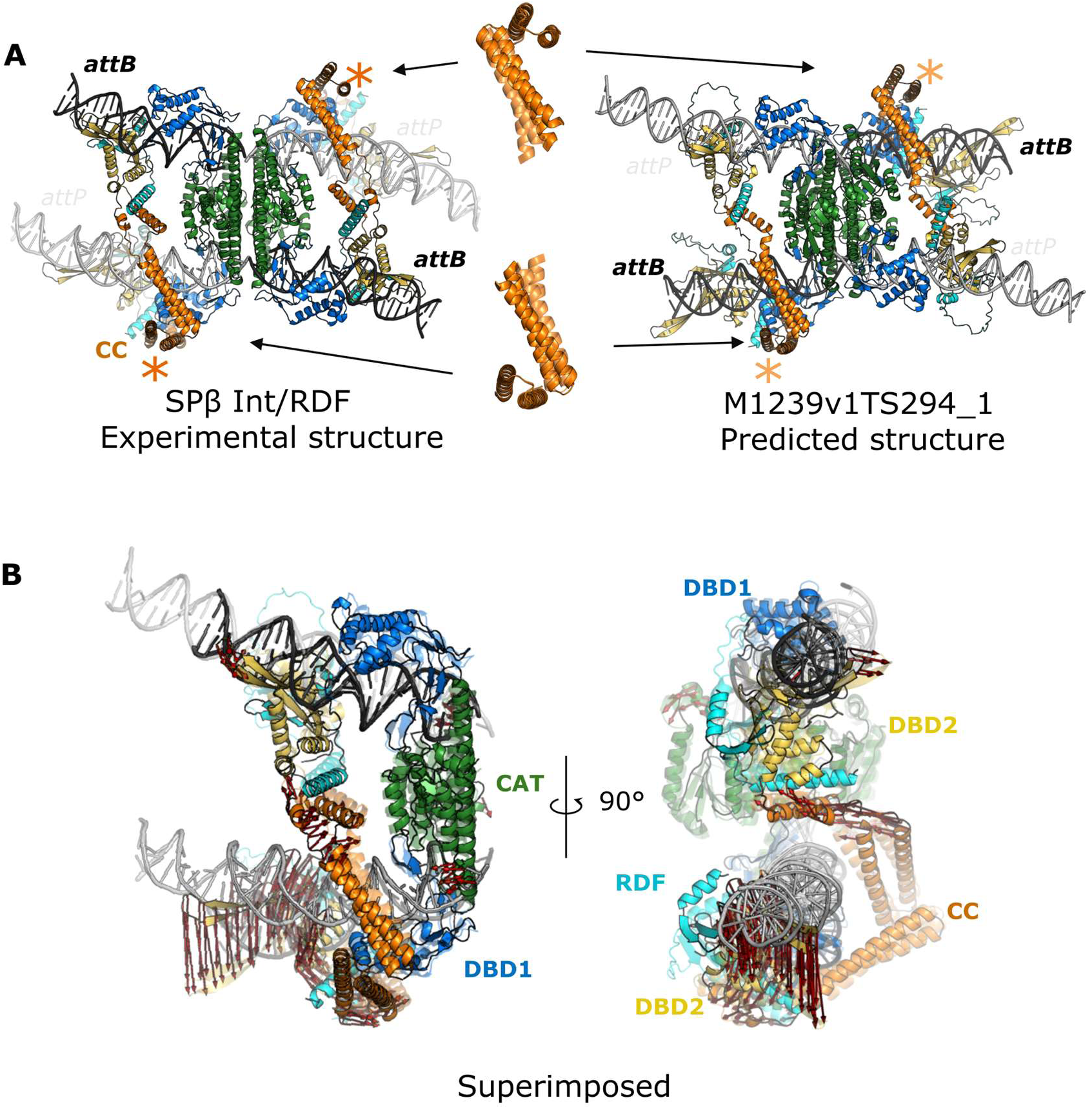
Experimental vs. predicted structure of SPβ LSI-RDF-DNA complex vs. predicted structure (CASP: M1239v1, PDB: 9DXD). (**A**) Cryo-EM structure of target M1239, the SPβ LSI-RDF synaptic complex in the pre-rotation state, (left) showing four distinct domains: an N-terminal catalytic domain (CAT, green), DNA binding domains 1 and 2 (DBD1 and DBD2, blue and yellow respectively), a coiled-coil motif subdomain (CC, orange), and a recombination directionality factor (RDF, cyan). The best CASP prediction model (M1239v1TS294_1, right) is shown on the right. Superimposed CC dimers from the experimental structure (dark orange) and the predicted structure (light orange) are aligned according to the CC-motif (residues 453-464), indicated by the asterisks (middle). (**B**) A superimposed half-complex of the experimental and predicted structures highlighting atomic distance (≥7Å) between the protein frameworks (C⍺) marked by red arrows, from the experimental to the predicted structure (transparency=0.5).

The overall scaffold of the synaptic complex consists of two *attL*-bound, chemically identical LSI-RDF dimers largely stabilized via hydrophobic interfaces among the N-terminal catalytic domains. In the center of this tetramer is a flat and hydrophobic platform where subunit rotation between left and right halves of the synaptic complex takes place during DNA recombination. Overall, this central tetrameric scaffold is very similar to that seen in 2005 for an otherwise quite different “small” serine recombinase^95^. In comparing 41 predicted models (M1239 category) to the SPβ LSI-RDF synaptic complex, 35 out of 41 entries predicted the overall tetrameric scaffold similar to the reference structure.

Overall, some but not all CASP models successfully predicted that there would be two different spacings of the two C-terminal DNA binding domains along the DNA. Most if not all successfully docked the DNA binding motifs with respect to the minor or major grooves of the DNA (**Figure 10A** right). However, the placement of the individual DNA binding domains did not correlate with the DNA sequence.

Nine out of the 35 entries with tetrameric cores were predicted to have CC-motifs engaged in a handshake interaction, and these entries are divided into two distinct modes: intra- and inter-dimer interactions (inter- is the correct configuration for this particular complex). Specifically, four entries (M1239v1TS033_1, M1239v1TS091_1, M1239v1TS110_1, and M1239v1TS167_1) adopted an intra-dimer handshake between CC-motifs. This interaction may be relevant to a post-DNA strand exchange conformation but is incorrect for this intermediate-state tetramer. Five entries (M1239v1TS231_1, M1239v1TS262_1, M1239v2TS262_1, M1239v1TS294_1, and M1239v2TS489_1) showed inter-dimer handshake interactions stabilizing synapsis between two DNA substrates, with the latter 4 of those resembling the overall scaffold seen experimentally, and with the entry from the Kihara lab (**Figure 10A** right) being the closest. However, none of the models matched the experimentally seen binding angle between the two CC-motifs (**Figure 10A**), perhaps because they also failed to bend the *attP* half sites toward the core tetramer rather than away as we see for *attB* half sites and in previously determined structures of distantly related small serine recombinases. The predicted *attP* bend may, however, reflect a pre- or post-cleavage state that we do not yet have an experimental structure for. Finally, none of the entries correctly modeled all 4 copies of the phosphoserine linkage between Ser22 and the 5’ phosphate of the DNA.

In conclusion, numerous entries successfully demonstrated that predicting protein structural asymmetry from chemically identical molecules within a synaptic complex was possible, and some of the models were, overall, remarkably similar to the experimental structure. However, sequence specificity in protein-DNA binding as well as atomic details for modeling the bending, compression, or stretching of DNA under tension—particularly generated from the covalently attached protein-DNA complex and from constraints in the CC-motif interactions—were significantly different from the experimental structure.

### 2.12 Cryptic DNA-binding protein UDE from *Cerapachys biroi* in complex with DNA (CASP: M1276) provided by Reinhard Albrecht, Yimin Hu and Marcus D. Hartmann

Due to a long-standing interest in uracil-binding proteins^96^, several years ago, our lab began investigating UDE, a protein implicated in the development of pupating insects (holometabola) in the late larval stages as a uracil-DNA degrading factor. First identified in *Drosophila*, this protein is primarily found in holometabola^97^, but also in a range of plant-pathogenic fungi. Canonical UDE is an all-helical protein with either two or three domains, comprising one or two copies of an N-terminal three-helix bundle domain, and a C-terminal domain consisting of six helices. In several species, it is also found as part of larger multi-domain proteins. Over the years, we have solved crystal structures of several two-domain UDE constructs from different species, including the yellow fever mosquito *Aedes aegypti* (*Aa*UDE; AAEL003864), which we had already entered as a target for CASP12. At that time, UDE constituted a new protein fold with no homologous structures available, and none of the participating groups was able to predict the entire UDE structure^98^. Today, however, very accurate UDE models are available in the AlphaFold database for many species, even though we have not yet released our experimental UDE structures.

After having solved the UDE structure, the most interesting aspect remaining was its DNA binding mode. To this end, we confirmed single- (ss) and double-stranded (ds)DNA binding activity for all constructs we could obtain for several species, as long as these constructs included at least one of the two possible N-terminal three-helix bundle domains. We found that this interaction was seemingly independent of the DNA sequence, and for some of the constructs, we were also able to obtain co-crystal structures with DNA fragments. Here, the highest resolution structure was obtained for a two-domain UDE construct from *Cerapachys biroi* (*Cb*UDE, A0A026WM60), the clonal raider ant, in complex with a ssDNA fragment, which we now entered as a hybrid prediction target into CASP16.

As anticipated from the binding experiments, the crystal structure revealed that the binding is mediated via the DNA backbone, without prominent interactions between the bases and the protein, and with an interface that is mostly involving the N-terminal domain. Of the 33 nucleotides of the ssDNA fragment used in the crystallization trial, only a stretch of about 8 nucleotides is well-resolved in the electron density. In the 5’ to 3’ direction, the first four of them are bound along the interface between the N- and C-terminal domain in a somewhat linear fashion, while the remaining four nucleotides are bound along the tip of the N-terminal domain (**Figure 11A**). In both segments, the bases protrude with a spacing resembling that in dsDNA. However, most interestingly, the two segments are separated by the intercalation of an arginine side chain located in the middle of the second helix of the N-terminal domain. This intercalation leads to a kink in the overall DNA trajectory and the geometry of the phosphodiester bond between the two segments. Although the functional and mechanistic details of UDE still remain to be uncovered, we expect this conserved arginine and its intercalation mode to play a key role.

**Figure 11.**
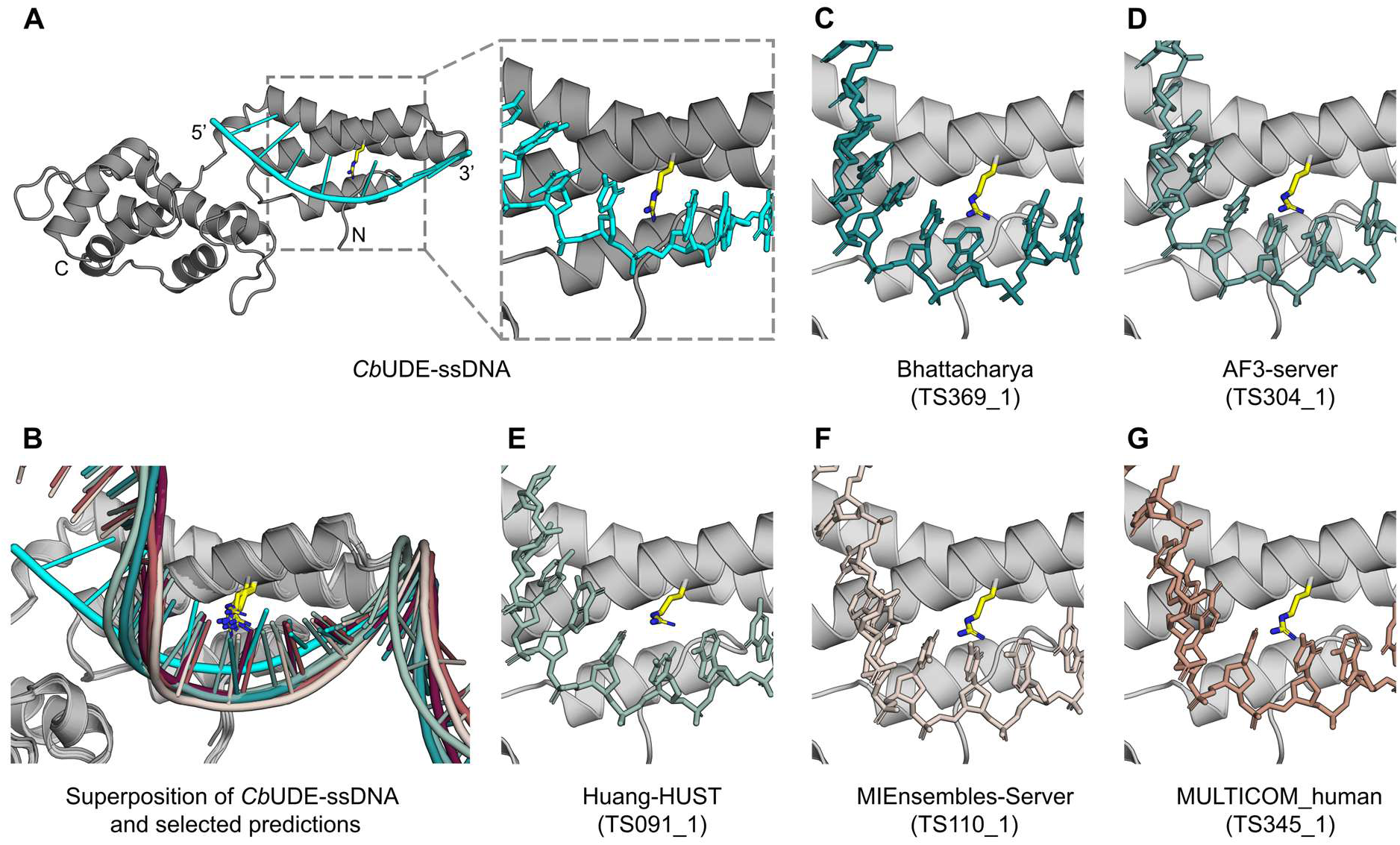
The crystal structure of *Cb*UDE in complex with ssDNA compared with the best first models submitted by the CASP16 predictors (CASP: M1276). (**A**) Crystal structure of *Cb*UDE bound to ssDNA. (**B**) Superposition of the crystal structure and the top eight first models as described in the text, including (**C**) Bhattacharya (TS369_1), (**D**) AF3-server (TS304_1), (**E**) Huang-HUST (TS091_1), (**F**) MIEnsembles-Server (TS110_1), and (**G**) MULTICOM_human (TS345_1), as well as MULTICOM (TS051_1), NKRNA-s (TS028_1), and Zheng (TS462_1).

While most of the groups participating in the hybrid prediction category in CASP16 did a good job of predicting the *Cb*UDE protein structure itself, only a few managed to predict relevant features of the DNA recognition mode. As judged by the first model submitted and in descending order, the best predictions are from Bhattacharya (TS369), AF3-server (TS304), Huang-HUST (TS091), MIEnsembles-Server (TS110), followed by identical models from MULTICOM_human (TS345) and MULTICOM (TS051), and essentially identical models from NKRNA-s (TS028) and Zheng (TS462), with Interface Contact Scores (ICS) and Interface Contacts Recall (ICR) scores above 0.5. When extending the evaluation to all five models submitted, also elofsson (TS241) is found in this leading cohort with a model identical to the first model of the AF3-server. While all these groups managed to predict the binding interface along the tip of the N-terminal domain, none of them predicted the intercalation of the arginine and the resultant kink in the DNA chain (**Figure 11B**). Rather, the DNA is modeled as a helical extension to the 3’ fragment bound to the N-terminal domain, clashing with the arginine, and consequently, none of the models predicted the interface responsible for the binding of the 5’ segment. In conclusion, while several groups were able to predict a part of the protein-nucleic-acid interaction site, further progress is needed to predict presumably functionally relevant details as those missed in the models for *Cb*UDE-ssDNA.

## 3 Conclusions

With the advent of accurate protein structure prediction algorithms, the nucleic acid community is eager for similar advances. Such algorithms should aim to predict the biologically, chemically, and biophysically important aspects of nucleic-acid containing structures. Through 12 examples, we have outlined the vital features of nucleic acid structures exposing areas for improvement in prediction accuracy. Critically, the detailed analysis by experimentalists reveals inaccuracies that CASP metrics may conceal, such as the enzymatically important η-η’ and π-π’ interactions of the group IIC intron and the binding site of SAMURI.

In most targets, secondary structure was predicted accurately, even for large targets with no homologous structures such as ROOL and OLE. However, multiple experimentalists note an over reliance on simple helical secondary structure units. For example, many important backbone kink, turns, or bends were not predicted accurately including the L2 bend in xrRNA, G-C domain in 3’ CITE, the target adenosine in SAMURI, the DNA near the intercalating arginine in UDE, and the DNA under tension in the SPβ LSI-RDF synaptic complex.

Further, non canonical interactions were underpredicted or poorly predicted in many targets such as xrRNA, RRE SLII, SAMURI, and the DNA aptamer. Some non canonical motifs and backbone kinks were predicted correctly, such as the kink turn in OLE, which was predicted correctly despite a contrary hypothesis in the literature. Additionally, there is some indication that evolutionary information may have assisted the prediction of tertiary interactions such as those in ROOL, even though global fold remained out of reach. Determining the global fold, or the arrangement of helices, remains a challenge more generally. For example, the orientation of the three helices of RRE SLII were predicted well when the junction was mutated to be simple (in target R1296) but very poorly when there were non canonical interactions at the junction (target R1209).

These non canonical interactions play a vital role in forming functionally important regions in nucleic acid structures. For example, they are integral to forming the active site and ligand binding site in SAMURI. We emphasize the importance of co-predicting the nucleic-acid structure with essential ligands. The nucleic-acid conformation is often changed when the ligand is bound. For example, dopamine intercalates between two bases in the DNA aptamer, the ZTP riboswitch changed conformation when a bulkier ligand was bound, and ions are vital for active site stability in group IIC intron. Such ligands are intrinsic structural elements and should be considered when modeling the nucleic acid structure. However, as noted for the DNA aptamer, many ligand predictions lacked basic chemical interactions integral to nucleic-acid-ligand interactions such as stacking and hydrogen bonding. This exposes nucleic-acid-ligand interactions as an area for improvement.

The principles considered in nucleic-acid-ligand interactions extend to RNA-RNA interactions and nucleic-acid-protein interactions. In both cases interfaces were generally underpredicted or predicted inaccurately. RNA multimer predictions were of poor quality, despite inter-molecular RNA-RNA contacts consisting of the same building blocks, such as base-pairs, as RNA tertiary structure. This suggests that knowledge of RNA tertiary structure was not directly transferable to RNA-RNA interface prediction. The novelty of RNA homo-multimerization as a prediction challenge was evidenced by predictors submitting asymmetric assemblies for ROOL, inaccurate or underpredicted interfaces for OLE, and dimers with no interface for the 2’F RNA origami. With the lack of previous examples, this poor performance is not surprising. Hopefully, the discovery of these complex RNA-multimeric structures will result in increased interface prediction accuracy in future challenges.

While there is more data on nucleic-acid-protein complexes, their predictions remain challenging except for known Fab-RNA interactions such as those used to engineer a crystallization construct for 3’CITE. Generally, the location of the interface on the protein was predicted accurately, but the interactions, such as the intercalation of an arginine of UDE into ssDNA, were not predicted accurately. For example, while the DNA helix was docked in the correct location of the SPβ LSI-RDF synaptic complex, all predictions placed the incorrect section of the DNA helix in the binding pocket. Finally, the majority of predictions placed the RNA aptamer in the incorrect location of the SARS-CoV 2 spike protein and those that did place it correctly, did not predict interactions accurately. The inaccuracy of the SARS-CoV 2 spike protein-RNA aptamer interface may be due to the 2’F modification of the RNA which predictors did not appear to consider. This modification also induced a dimerization of an RNA origami which does not dimerize without modification, highlighting the importance of considering nucleotide modifications. In the future, the CASP target definition format might be expanded to provide a machine-readable field or format for nucleotide modifications.

We note that predictions that deviate from the experimental structure may be valid biologically-relevant structures. The experimental structure presented here-in do not represent the only ground-truth structures – they represent the structure(s) of the molecule under specific experimental conditions and time points. For example, predictions for the SPβ LSI-RDF synaptic complex could represent different states along the catalytic cycle of recombination or predictions of the RRE SLII could represent different states in the solution ensemble of structures it occupies that were not crystallized. This offers challenges and opportunities for the structure prediction community including how to consider experimental conditions in structure prediction and how to assess the accuracy of both the predictions and experimental structures, perhaps by systematic comparison to independently collected evolutionary and experimental data. In the future, experimentalists could be asked to contribute data containing information about the ensemble of structures, such as particle stacks of cryo-EM data, to tackle these challenges.

Overall, we observe accurate predictions of nucleic acid secondary structure and in some cases global folds, but warn against over-reliance on canonical structural motifs, in particular in functionally important regions. We urge improvement in the prediction of backbone kinks and non canonical base-pairing. The prediction of nucleic-acid interactions with proteins, ligands, and other nucleic acids should remain emphasized in future CASPs to drive progress in this new frontier for structure prediction. Due to the effort of experimentalists across the world, CASP16 presented nucleic acid structures with novel folds, novel interactions, and even novel prediction categories, demonstrating that there remain underexplored aspects of nucleic-acid structure only unveiled by future experimental structure determination.

## Author Contributions

Concept, abstract, introduction, conclusion, editing, and coordination: R.C.K, A.K., and J.M.. Target-specific sections: by authors provided in the sections’ titles.

## Acknowledgments

We acknowledge the following sources of funding and support: US National Institute of General Medical Sciences (R01GM100482 to A.K., R35GM149336 to J.A.P., Northeastern Collaborative Access Team beamlines P30 GM124165); National Institutes of Health grants (R01GM079429 to W.C., R35GM122579 to R.D., R35GM150778 to A-L.S, T32 grant GM066706 to M.O., transition award K22HL139920-01 to C.P.J., intramural program of the National Heart, Lung and Blood Institute); National Institutes of Health Common Fund Transformative High-Resolution Cryo-Electron Microscopy program (U24GM129541); Center for Structural Biology of HIV-1 RNA (CRNA) Collaborative Development Pilot Grant Program (NIH NIAID1U54AI170660, Sub-award SUBK00019302) to D. K.; the National Science Foundation (2330652 to W.C. and R.D., GRFP DGE-2036197 to J.G.G.); Bio-X Bowes Graduate Student Fellowship (R.C.K.); the Howard Hughes Medical Institute (to R.D.); National Key Research and Development Program of China (2022YFC2303700 to Z.S.); Natural Science Foundation of China (32222040 to Z.S.); Swedish National Research Council (VR, 2024-04107 to M.M.); HORIZON-MSCA-2023-DN-01 action (project: TargetRNA, n. 101168667 to M.M.); France Canada Research Fund (to C.D.M. and P.E.J.); Natural Sciences and Engineering Research Council of Canada (to P.E.J.); scientists of the EMBL Grenoble; scientists of the EMBL Heidelberg; ESRF CM01 cryo-EM facilities; Thomas Mulvaney, Mauro Maiorca and Maya Topf at CSSB Hamburg; EMBION Cryo-EM Facility at iNANO, Aarhus University; Novo Nordisk Foundation (NNF21OC0070452 to E.S.A); the Villum Foundation (VIL71957 E.L.K); Andrei Lupas; beamline X10SA/Swiss Light Source; Eiger 16M detector on 24-ID-E is funded by a NIH-ORIP HEI grant (S10OD021527); Advanced Photon Source, a U.S. Department of Energy (DOE) Office of Science User Facility operated for the DOE Office of Science by Argonne National Laboratory under Contract No. DE-AC02-06CH11357; Deutsche Forschungsgemeinschaft (project no 463143961 and Gottfried Wilhelm Leibniz Programme to C.H.). This article is subject to HHMI’s Open Access to Publications policy. HHMI lab heads have previously granted a nonexclusive CC BY 4.0 license to the public and a sublicensable license to HHMI in their research articles. Pursuant to those licenses, the author-accepted manuscript of this article can be made freely available under a CC BY 4.0 license immediately upon publication.

## Data Availability

The data that support the findings of this study are openly available at the CASP website, the scores can be found at:

https://predictioncenter.org/casp16/results.cgi?tr_type=rna

https://predictioncenter.org/casp16/results.cgi?tr_type=rna_multi

https://predictioncenter.org/casp16/results.cgi?tr_type=hybrid

The predicted models can be found at: https://predictioncenter.org/download_area/CASP16/predictions/RNA/

https://predictioncenter.org/download_area/CASP16/predictions/oligo/

https://predictioncenter.org/download_area/CASP16/predictions/hybrid/

